# A pan-MHC reference graph with 246 fully contiguous phased sequences

**DOI:** 10.1101/2023.09.01.555813

**Authors:** Liza Huijse, Solomon M. Adams, Joshua N. Burton, Julianne K. David, Russell S. Julian, Galit Meshulam-Simon, Harry Mickalide, Bersabeh D. Tafesse, Verónica Calonga-Solís, Ivan Rodrigo Wolf, Ashby J. Morrison, Danillo G. Augusto, Solomon Endlich

## Abstract

The major histocompatibility complex (MHC) is a region of the human genome that is key to immune system function but sometimes refractory to genomic analyses due to extreme polymorphism and structural variation. We performed targeted long-read sequencing and *de novo* assembly of MHC to create 246 highly accurate, fully contiguous, and phased full-length sequences, mostly from data provided by the Human Pangenome Reference Consortium (HPRC). We identified alleles at high resolution across 39 loci including the class I and II HLA (human leukocyte antigen) genes, discovering 1,246 putative novel allele sequences. We identified copy number variation in the *C4A* and *C4B* genes and found significant linkage disequilibrium between *C4A∼C4B* haplotypes and 14 MHC loci. We build our sequences into a novel “pan-MHC” reference graph, and we demonstrate that this improves the accuracy of short-read variant calling. Our haplotypes and graph contain significantly more population diversity than preexisting MHC sequences, thus improving the prospects for global health equity in this clinically important genomic region.

## Introduction

The major histocompatibility complex (MHC) is a gene-dense region on chromosome 6 of the human genome, with a length that is 3-5 Mb, depending on its definition, but varies among individuals^1–4^. It encompasses approximately 165 protein-coding genes, many of which are critical to immune system function, most notably the human leukocyte antigen (HLA) gene complex^2^. The main biological function of HLA genes is related to antigen presentation of self- and non-self-peptides to T cells^5,6^. HLA genes were first discovered via their central role in transplantation^7^ and have also long been appreciated for their role in pharmacogenomics^8^. More recently, genome-wide association studies (GWAS) have identified far more trait associations with the MHC than with any other genomic region, confirming its exceptional functional significance^9–11^. Thus MHC, including HLA, is an extremely promising candidate region for disease studies, biomarker discovery, and personalized medicine.

Despite its clinical importance, accurate and broad characterization of MHC remains out of reach for most clinicians and researchers. Genes in the MHC region have high levels of sequence variation, structural variation (SV), and homology. Moreover, some genes feature extreme linkage disequilibrium (LD)^12^, which confounds the identification of causal variants^13,14^. The MHC region also contains many kinds of repetitive elements^15^, posing challenges to assembly and alignment methods. Most significantly, the MHC region is the most polymorphic part of the human genome, with an enormous number of alleles known and likely many more yet to be discovered, such that no single reference sequence comes close to properly representing it^16,17^. A particularly intractable area of the MHC is the one hosting *C4A* and *C4B* genes, which encode for C4 (complement component 4) proteins. *C4A* and *C4B* are clinically relevant^18–20^ and are known for their extensive copy number variation (CNV) and high levels of sequence identity^21^. The common practice of using static linear reference genomes creates reference bias in variant and allele calling for genes within the MHC region^22^. Reference bias is especially pronounced when studying non-European populations who are historically underrepresented in medical research and is thus a threat to health equity^23^.

A comprehensive workflow for MHC genomic analysis would entail calling all alleles of all genes, phasing all alleles, discovering novel alleles if present, and identifying CNV in *C4A* and *C4B*, all at high enough throughput to handle large numbers of samples^24^. It should also identify non-coding variants, which play a major role in HLA gene regulation^14,25^. HLA genes, for instance, are generally genotyped by commercially available kits that do not always include all genes or achieve maximum resolution^26^. The IMGT/HLA database^17^, a public repository for known HLA allele sequences, can be used as a reference for HLA genotyping; however, a pre-existent database of alleles might not always allow the discovery of new alleles, nor can it easily identify haplotypes or intergenic variation. Custom bioinformatic methods exist to discover novel HLA alleles using short reads^27,28^, but throughput and phasing remain challenging. To solve the problem of phasing, dense genotyping arrays have been used to construct large population-specific reference panels and quantify LD^29,30^, but this approach is expensive and only applicable to one population at a time.

Long-read approaches to MHC analysis exploit the recent availability of long sequencing reads, such as from PacBio and Oxford Nanopore platforms. These reads are long enough to span entire HLA genes, capturing coding and non-coding variation and yielding phased full-length sequences of novel alleles^31^. They can even, with the aid of specialized bioinformatic methods, create complete, phased haploid MHC sequences^32,33^. Fully phased MHC sequences are invaluable resources because they can serve as references for imputation using short reads. Very few of these sequences exist today, to our knowledge, and they are mostly derived from individuals of European ancestry^32^, so are blinded to the great diversity of HLA alleles and haplotypes in non-European populations^25,34–37^.

Pangenome reference graphs are gaining popularity as a method for systematically handling the diversity of human genomes^23,38–40^, and because no genomic region is more diverse than MHC, it is in particular need of a graph. Reference graphs are computational structures that encode multiple sequences in order to represent variation within a population^41^. Compared to traditional linear reference sequences, graphs provide superior read mapping and variant calling by reducing reference bias^23,38,41^. Many graph-based bioinformatics toolkits have been developed in recent years, including vg^38^, GRAF^42^, and ODGI^43^. Reference graph methods have also been optimized for the MHC region^44,45^ but these graphs, like other MHC resources, are limited by the small number of known MHC sequences. New methods are needed both to expand the scope of MHC region knowledge and to translate that knowledge into tools for variant and allele calling.

Here we use long-read sequencing data to assemble complete, phased MHC sequences and construct an MHC reference graph. We apply our assembly method to 119 samples from the Human Pangenome Reference Consortium (HPRC)^39,40^ for which long reads are publicly available, as well as to four cell lines we processed using our novel wet-lab method for target enrichment and sequencing. In total, we assemble 246 fully phased MHC sequences, to our knowledge the largest collection to date, and we discover 1,246 alleles that are absent from the IMGT/HLA database. We integrate these haplotypes into a “pan-MHC” reference graph spanning diverse human populations, and we demonstrate the superiority of this graph for calling variants in MHC from short-read sequencing data.

## Results

### We created 246 highly accurate, fully contiguous, and phased full-length MHC sequences

We sequenced the cell lines DBB, QBL, PGF, and HG00733 on a PacBio Sequel II instrument, using both whole-genome sequencing and targeted sequencing of MHC. For these samples, our targeted sequencing protocol achieved enrichment rates of as high as 94×, though rates of around 30× were more common (**Supp. Fig. 1**), and average read lengths between 8.9 kb and 15.7 kb. We then used these readsets to create fully contiguous assemblies of the MHC region (**Fig. 1**). Additionally, we downloaded 130 publicly available datasets of >30× PacBio coverage depth of cell lines sequenced by the HPRC^40^ and assembled their MHC regions (see **Methods**). Of these 130 samples, 11 (8.5%) failed to create fully contiguous diploid MHC assemblies, while the remaining 119 samples were assembled successfully and were used in further analysis (**Supp. Table 1**). These 119 samples plus the DBB, QBL, PGF, and HG00733 assemblies made a total of 123 *de novo* assemblies, and thus a total of 123 × 2 = 246 contiguous phased sequences (**Table 1**). We refer to these sequences as *haplotigs*. Nearly all haplotigs had notable local and structural variation relative to the reference MHC sequence in the GRCh38 reference genome (**Fig. 2**).

**Figure 1:**
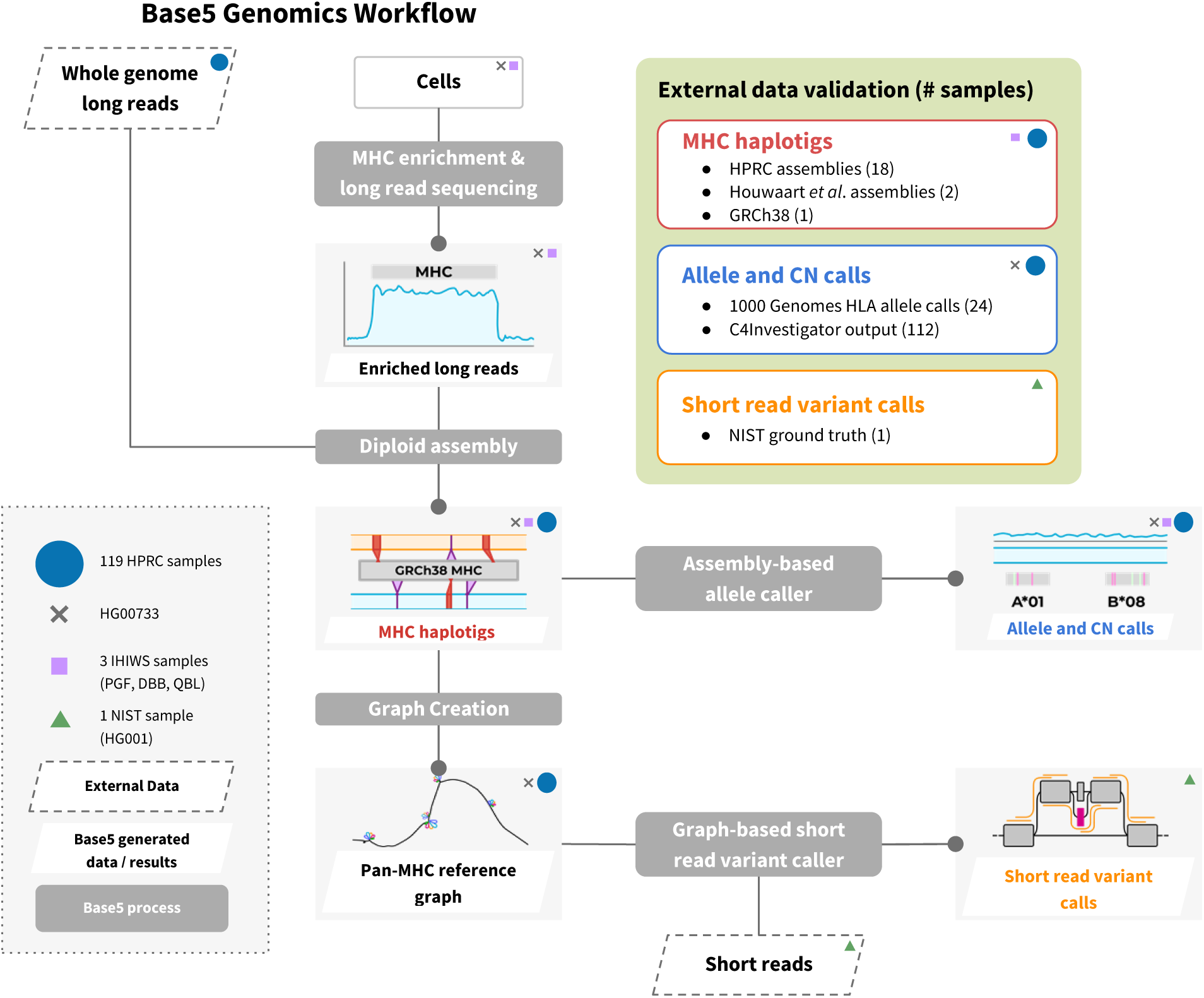
Overview of experimental setup. We subjected four cell lines to targeted enrichment of the MHC region and PacBio HiFi sequencing. Additionally, we downloaded whole-genome PacBio HiFi reads from 119 cell lines generated by the HPRC^40^ and selected their MHC-region reads. We assembled each of these 123 samples’ MHC-targeted readsets into diploid, fully sequence-contiguous MHC haplotigs. We called the alleles and copy numbers of MHC loci directly from the assembled sequence. We also built the haplotigs into our flagship pan-MHC graph, and we leveraged this extensive graph for use in a graph-based short-read variant calling tool. Validation methods included: internal quality controls; comparison of 21 of the cell lines’ assemblies with GRCh38 and with published assemblies from HPRC and Houwaart *et al*.^32^; comparison of 24 samples’ HLA allele calls and 112 samples’ C4 copy number calls with output from state-of-the-art external tools^37,52^; and comparison of short-read variant calls for 1 sample with ground truth variant calls published by NIST.

**Figure 2:**
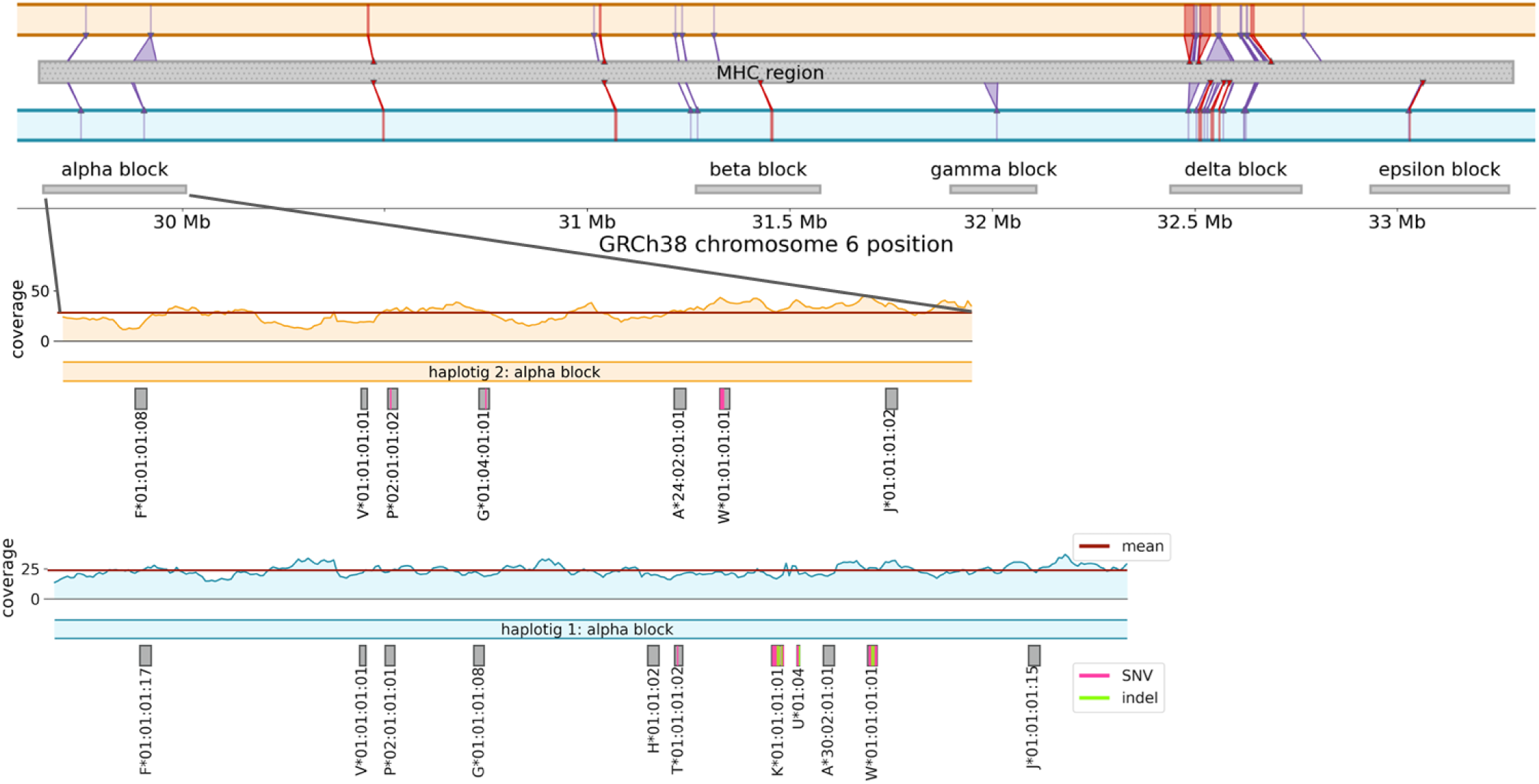
Sequence contiguity of haplotigs and their variance from reference and each other. The upper panel shows our two contiguous HG00733 haplotigs 1 (blue, lower bar) and 2 (orange, upper bar) and their large structural variation against the GRCh38 reference sequence. Purple and red wedges show regions of large (≥1kb) deletions and insertions, respectively, from the reference. The lower two panels expand the region of the alpha polymorphic frozen block for haplotigs 1 (lower, blue) and 2 (upper, orange), where the upper panel segment shows HiFi read depth across the region, with mean depth in maroon. The identified gene locations and their called alleles are shown below, with locations of point mismatches and indels between the haplotig sequence and the IMGT/HLA^17^ allele database sequence marked in pink and green, respectively.

**Table 1:** Summary statistics of haplotigs and graphs.

| Statistic | MHC | HOXA |
| --- | --- | --- |
| Number of haplotigs input to graph | 240 | 230 |
| Total length of haplotigs input | 844 Mb | 997 Mb |
| Mean length of haplotigs input | 3.52 Mb | 4.33 Mb |
| Bp alt sequence per 1 Mb input sequence | 12,227 | 93 |
| N nodes in graph, per 1 Mb input sequence | 123 | 48 |
| N long nodes ($\geq 50$ bp) in graph, per 1 Mb input sequence | 2.0 | 0.1 |
| GC content of alt sequence | 0.424 | 0.432 |
| Number of loci at which alleles are called | 39 | N/A |
| Number of total observed instances of these loci | 8,746 | N/A |
| Number of observed instances of novel alleles | 2,580 | N/A |
| Number of distinct novel allele sequences | 1,246 | N/A |

### Haplotigs have a mean phase contiguity-50 of 1.66 Mbp, and 85% have no mismatches with read alignments

Our ability to determine haplotype phase across the MHC region is limited by the length of our long reads and, relatedly, by long regions of homozygosity in an individual. To evaluate our assemblies’ phasing, we aligned each sample’s HiFi reads to the sample’s two haplotig sequences, where a relative drop in coverage depth of unambiguously aligning reads indicates a potential loss in phase contiguity (defined as contiguity of phasing without switch errors). We found that 83.6% of all assembled PFBs (polymorphic frozen blocks, regions of relatively high diversity and low recombination^15,46^) were fully phase-contiguous, 50.8% of assembled haplotigs (130/246) had all PFBs completely phased, and 6.9% (17/246) had complete phase contiguity across the entire haplotig (**Supp. Fig. 2a**). Haplotigs had a median of 4 regions of potential phase loss each, with mean and median phase loss region lengths of 48,028 and 6,559 bp respectively (**Supp. Fig. 2c**). To quantify the phase contiguity of our haplotigs, we define a new metric, phase contiguity-50 (PC50), analogous to N50 for sequence contiguity. PC50 is defined as the length *X* at which at least half the haplotigs’ bases are in phase contiguity with ≥*X* bases (see **Methods** for details). The mean of haplotig PC50 values across all 246 haplotigs was 1.66 Mbp (megabase pairs) (**Supp. Fig. 2b**).

We reasoned that phase loss was prone to occur within long stretches of homozygosity. We aligned each sample’s two haplotigs to each other and found that regions of potential phase loss indeed correlated with homozygous stretches: 73% of all phase loss regions and 94% of phase loss regions >5kb overlapped with a homozygous region >10kb (**Supp. Fig. 3**). Therefore nearly all of our breaks in phase contiguity are attributable to underlying MHC homozygosity.

Our read alignments to haplotig sequences also allowed us to investigate our haplotigs for local misassemblies. For 84.6% of haplotigs (208/246), the high-quality aligning reads show no systematic mismatches (SMs). Of the remaining 38 haplotigs, 26 have only a single such mismatch over the entire ∼3.5-Mb haplotig length, and only five haplotigs show evidence of significant misassembly (**Supp. Fig. 2d**). Four of these have over 10 SMs with a concentration of 0.013-1.2 SMs per kilobase of contiguous sequence, and the final haplotig shows evidence of a phase switch occurring in a heterozygous region.

### Haplotigs have >99.999% sequence identity with HPRC assemblies

We evaluated our assemblies for sequence and phasing accuracy by comparing them with whole-genome assemblies of some of the same samples recently published by HPRC^40^. The MHC region (when defined as GRCh38 region chr6:28,510,120-33,480,577) was found to be contiguous in 54.3% (25/46) of the HPRC diploid assemblies, compared to 89.2% (116/130) of our assemblies. We compared 18 diploid assembly pairs (those that could be matched with HPRC) in detail and observed a very high degree of sequence level identity (**Fig. 3a**). On average we observed 4 mismatches, including 3.6 gaps and 0.3 substitutions, per million bases, between matched haplotigs from HPRC and the current study - a sequence identity of 99.9996%. The largest discrepancy was regarding phasing, for which we observe an average of 0.1 phase switches per million bases. When we accounted for known loss of phase across long homozygous regions (regions larger than the average read length (**Supp. Fig. 4**)), we found 5 remaining phase switches in >125 million total bases that were true discordances.

**Figure 3:**
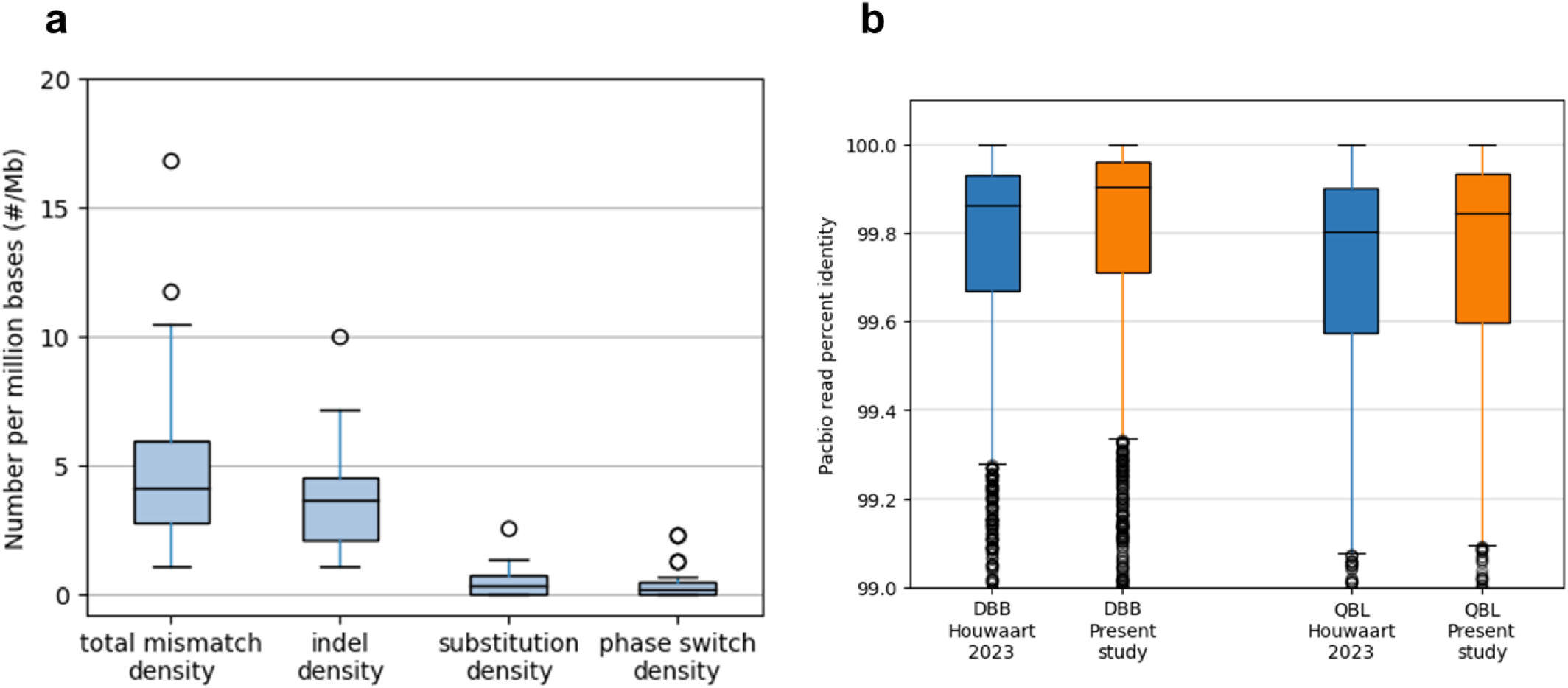
Assembly sequence validation. **a.** Comparison of our assemblies to the MHC regions of whole-genome assemblies from HPRC^40^, performed by haplotig-to-haplotig alignment of 18 matched samples. Total mismatches includes both substitutions as well as insertions and deletions (indels). Indels are counted in units of gap openings. Phase switches are identified as regions where the sequence of our haplotig 1 switches from resembling HPRC’s haplotig 1 better to resembling HPRC’s haplotig 2 better and vice versa. Apart from phase switches, we did not observe any structural differences. **b.** Comparison of the quality of our DBB and QBL assemblies to external assemblies from Houwaart *et al*.^32^, using alignments of PacBio reads. We took PacBio readsets not used in the creation of either assembly, and aligned them to each assembly using minimap2^65^. To highlight differences between the assemblies, we retained only alignments which overlapped positions where our assembly differs from the external one. *y*-axis: percent identity of the alignments.

We furthermore compared inspection of raw read alignments to our haplotigs and the HPRC haplotigs for the 18 matched samples. Nine of our haplotigs have SMs between reads and assemblies while eight HPRC haplotigs have similar SMs. For most haplotigs, the SMs counts are all very low (from 1-3 per haplotig for our assemblies and 1-2 for the HPRC assemblies), but one of the HPRC haplotigs shows evidence of significant misassembly (48 SMs over a sequence length of 17,672 bases) **(Supp. Fig. 5)**. Overall, this leads to a significant-misassembly rate of 2.0% (5/246 haplotigs) for our assemblies, compared with 2.8% (1/36 haplotigs) for the HPRC assemblies. In summary, raw read alignments demonstrate that the structural quality of our assemblies meets that of the HPRC’s MHC-region assemblies.

### Our assemblies of DBB, QBL, and PGF are the most accurate to date

The DBB, QBL, and PGF cell lines have had their MHC regions thoroughly characterized^16,32^, enabling us to perform an in-depth validation of our assembly performance. We compared our PGF assembly to the MHC region of GRCh38, which is derived from PGF^47^, and we compared our DBB and QBL assemblies to the assemblies generated by Houwaart *et al*.^32^. We replicated the known feature of high MHC homozygosity in these cell lines: sequence identity between the two haplotigs (throughout our defined MHC region) was 99.9997% for DBB, 99.9868% for QBL, and 99.9999% for PGF. Our assemblies also had high sequence identities with the existing assemblies: for PGF our assembly has 99.996% sequence identity with the MHC region of GRCh38; for DBB and QBL our assemblies have 99.97% sequence identity to their respective Houwaart *et al*. assemblies. To assess these differences between assemblies, we aligned external test readsets to them (see **Methods**). Using PacBio readsets of DBB and QBL, we found that the reads match our assemblies more closely than the Houwaart *et al*. assemblies (sequence divergence of 0.09% vs. 0.13% for DBB and 0.15% vs. 0.19% for QBL) (**Fig. 3b**). Using Illumina readsets of DBB and QBL, we obtained sufficient coverage depth to call variants with DeepVariant^48^, which found several times as many errors in the Houwaart *et al.* assemblies as in our assemblies: 438 vs. 77 for DBB and 422 vs. 58 for QBL (**Table 2**). We applied this same analysis to PGF and identified 75 errors in our assembly and 75 in GRCh38, suggesting similar quality. In sum, external readsets suggest that our MHC assemblies of these cell lines are of similar or higher quality than any other publicly available assemblies.

**Table 2:**
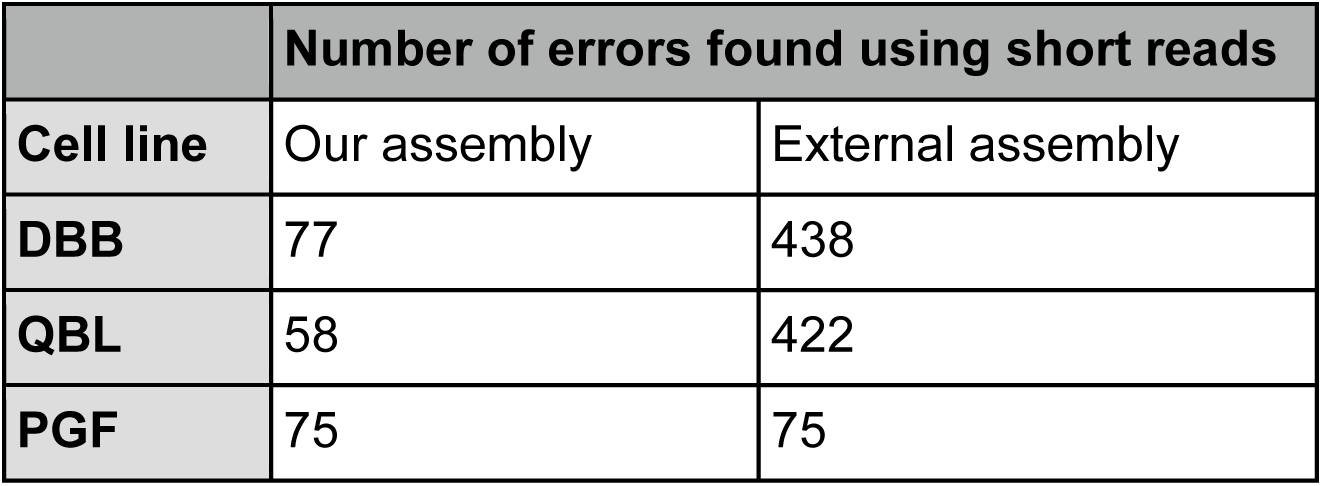
Estimates of assembly error rates from short-read alignments. We aligned short reads to all assemblies and called variants with DeepVariant^48^. We defined “errors” as variant calls that were either homozygous or compound heterozygous.

|  | Number of errors found using short reads |  |
| --- | --- | --- |
| Cell line | Our assembly | External assembly |
| DBB | 77 | 438 |
| QBL | 58 | 422 |
| PGF | 75 | 75 |

### Identifying and discovering HLA alleles at maximum resolution

Our fully contiguous MHC haplotigs contained all the necessary information to identify HLA alleles at maximum (four-field) resolution, which includes synonymous and noncoding variation; it also permits novel allele discovery and identification of phased haplotypes. We designed a custom tool to find allele sequences in the haplotigs and applied it to 39 loci in the IMGT/HLA database^17^. Across our 246 haplotigs, we made a total of 8,746 allele identifications (**Table 1**). Among these allele calls, 5,895 had sequences perfectly matching a sequence previously deposited in IMGT/HLA. Another 271 calls diverged from IMGT/HLA entries only by an A or T indel within a >10-bp homopolymer region; we discounted these sequences due to uncertainty in the accuracy of long homopolymers in PacBio sequencing. The remaining 2,580 allele calls are indicative of novel alleles (**Supp. Fig. 6**). To validate these novel allele calls, we performed a read alignment inspection (see **Methods**), which 2,570 (99.6%) passed (**Supp. Fig. 7**). We find substantial overlap in our novel allele calls: these 2,570 calls encompass only 1,246 unique sequences, which represent putative discoveries of alleles not known to the IMGT/HLA database (**Supp. Table 2**). Of these alleles, 494 (39.6%) involve coding sequence variation. Among the 11 particularly well-studied *HLA* loci covered by Thermo Fisher’s AllType kit^26^, we saw 271 putative novel allele sequences, of which 35 (12.9%) involve coding sequence variation (**Supp. Fig. 8**).

We compared our allele calls to a set of *HLA* allele calls made by the PolyPheMe software^37^. PolyPheMe uses exome sequencing to call alleles at 2-field resolution (coding variants only) at five *HLA* loci: *A*, *B*, *C*, *DRB1*, and *DQB1*. Abi-Rached *et al.* used PolyPheMe to identify HLA alleles in 24 of the same cell lines we studied. Our results were concordant with PolyPheMe’s at 2-field resolution for 228 of 230 allele calls that could be compared (99.1%) (**Supp. Table 3**).

### High-resolution insights into *C4A∼C4B* haplotype structure

Next, we sought to investigate the region that includes the *C4A* and *C4B* genes. These two genes play significant and functionally distinct roles^49–51^ yet share more than 99% sequence identity and exhibit extensive CNV across individuals and populations, which has confounded previous analyses^18,52^. At a sequence level, our haplotig assemblies contained 467 novel allele sequences in *C4A* and *C4B* (median 2 per haplotig). At a structural level, our haplotigs contained multiple copies of *C4A* and *C4B*, in long and/or short isoforms (designated -L, -S) (**Fig. 4a**). The minimum observed number of copies of *C4A* and/or *C4B* was one (28 haplotigs), while the maximum was five (1 haplotig). We observed the *C4A* and *C4B* genes in the standard structure configuration *C4A*-L∼*C4B*-L∼*C4A*-S∼*C4B*-S. For example, when *C4A*-S and *C4B*-S co-occurred on the same haplotype, *C4B*-S never preceded *C4A*-S, similar to previous observations^18^.

**Figure 4:**
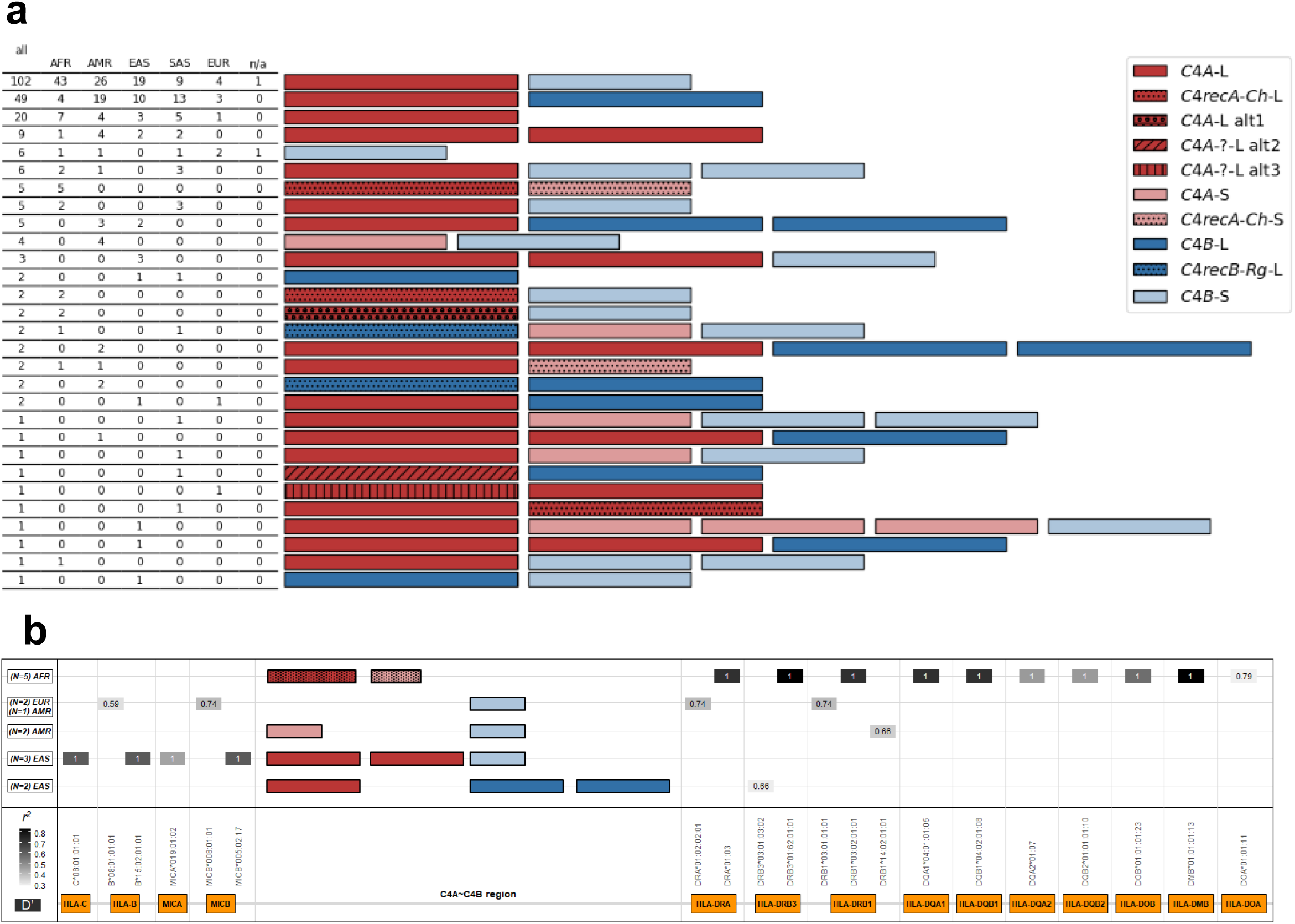
*C4A∼C4B* structure and CNVs in our assemblies. **a.** All C4 haplotype groups discovered in our 240 MHC haplotigs (PGF, QBL, and DBB were excluded in this analysis). *C4A* genes are shown in red and *C4B* in blue, with darker color indicating a long allele and lighter color indicating a short allele. Unshaded boxes indicate the expected blood group match, *i.e.*, Rodgers (Rg) for *C4A* and Chido (Ch) for *C4B*; dotted hatching indicates recombinant blood groups, and other hatch patterns indicate other variants. Novel alleles, which we name here, include alt1, which has a single variant in a *C4A*-defining base position, and alt2 and alt3, which have variants in the blood group-defining base positions. **b.** Linkage disequilibrium (LD) events involving C4. The bottom row with orange boxes shows genomic loci in the MHC region. The five upper bars show statistically significant LD events (*p* < 10^-14^) between C4 CNVs and the indicated allele (**Supp. Table 5**). Squares indicate LD *r^2^* values (grey shade) and *D’* values (number), and the allele names below the squares show the alleles in LD.

For each sample, we called sample-level copy number (CN) for C4 by adding up the observed CN of all *C4A* and *C4B* loci on both haplotigs. To validate these calls, we downloaded 30× coverage depth of Illumina sequencing data of 112 of our samples from the International Genome Sample Resource (IGSR)^53^ and called *C4A* and *C4B* CNs using C4Investigator^52^. C4Investigator’s CN calls were concordant with ours for 106 of 112 samples (94.6%), and all discordant calls were different by only 1 CN (**Supp. Table 4**). Thus our CN calls in the *C4A∼C4B* region successfully replicate read-depth-based methods.

Apart from the set of five SNPs that distinguish the primary function of C4A and C4B proteins, another set of four SNPs in *C4A* and *C4B* defines the blood groups Rodgers (Rg) and Chido (Ch). These two SNP regions are separated by only 456 bp and are in tight LD. As a result, typically proteins encoded by C4A are characterized by the presence of Rg, while C4B proteins carry Ch^52,54^. C4A and C4B loci encoding for proteins *C4recA-Ch* and *C4recB-Rg* are considered recombinant. These recombinants were recently observed at high frequencies, especially in Africans^52^. We looked for such recombinants in our assembled haplotigs. Among our 240 haplotigs (excluding DBB, QBL, and PGF) we observed 10 haplotigs (4.2%) containing *C4recA-Ch*; 4 (1.7%) containing *C4recB-Rg*; and 2 (0.8%) containing novel *C4A* sequences that could not be easily classified as Rg or Ch (**Fig. 4a**; **Table 3**). Most haplotypes carrying recombinant genes presented a non-recombinant gene in the same haplotype; the only exception was a haplotype consisting of two *C4recA-Ch*, which appeared in five individuals, all of West African ancestry (**Fig. 4a**).

**Table 3:** Counts of C4 allele types on our 240 haplotigs. The column “Total allele CN” tallies the total number of times that allele appears on any haplotig, including multiple appearances on one haplotig. The second column, “Number of haplotigs”, tallies the number of haplotigs on which the allele appears.

| Allele | Total allele CN | Number of haplotigs |
| --- | --- | --- |
| **C4A-L** | 229 | 213 |
| **C4B-L** | 73 | 66 |
| **C4B-S** | 145 | 137 |
| **C4A-S** | 11 | 9 |
| **C4recA-Ch-L** | 8 | 8 |
| **C4A-L alt1** | 2 | 2 |
| **C4recA-Ch-S** | 7 | 7 |
| **C4recB-Rg-L** | 4 | 4 |
| **C4A-?-L alt2** | 1 | 1 |
| **C4A-?-L alt3** | 1 | 1 |

To our knowledge, LD between *C4A∼C4B* haplotypes and *HLA* alleles has never been tested at high resolution. With phased data in hand, we found significant LD between five *C4A∼C4B* variants and alleles of 14 MHC loci (**Fig. 4b; Supp. Table 5**). Interestingly, the double-recombinant *C4recA-Ch∼C4recA-Ch* found in five West African individuals was in LD with *HLA-DRA*01:03* (*D’*= 1; *r^2^* = 0.71).

### A “pan-MHC” reference graph

We built our 240 MHC-region haplotigs into a “pan-MHC” reference graph, which, thanks to the diversity of HPRC, already represents more ancestral groups than previously reported (**Fig. 5a**; **Table 4**). The graph is able to produce base-resolved end-to-end alignment of our target regions, even in the most complex areas of C4 (**Supp. Fig. 9**). For comparison, we chose a control genomic region around *HOXA* (defined by the region chr7:25296538-29630479 on GRCh38) as a representative of a potentially non-polymorphic locus given its critical role in development and conservation among species^55^. As expected, we were able to confirm that this control region has a relatively low rate of small and structural variation relative to MHC. We assembled 230 sequence-contiguous haplotigs in this control region and built these into a “pan-HOXA” graph (**Fig. 5b**). The average MHC haplotig length (corresponding to the GRCh38 region chr6:29,668,442-33,184,109) was 3.50 Mb, with a coefficient of variance (CV) of 2.0%, and the pan-MHC graph contained 10.3 Mb of sequence not represented in GRCh38. In contrast, for the region around *HOXA*, the CV of haplotig size was only 0.08%, and the pan-HOXA graph contained only 92.3 kb of sequence not in GRCh38 (**Fig. 5c**; **Table 1**). This confirms the extreme polymorphism of MHC compared to the control region.

**Figure 5:**
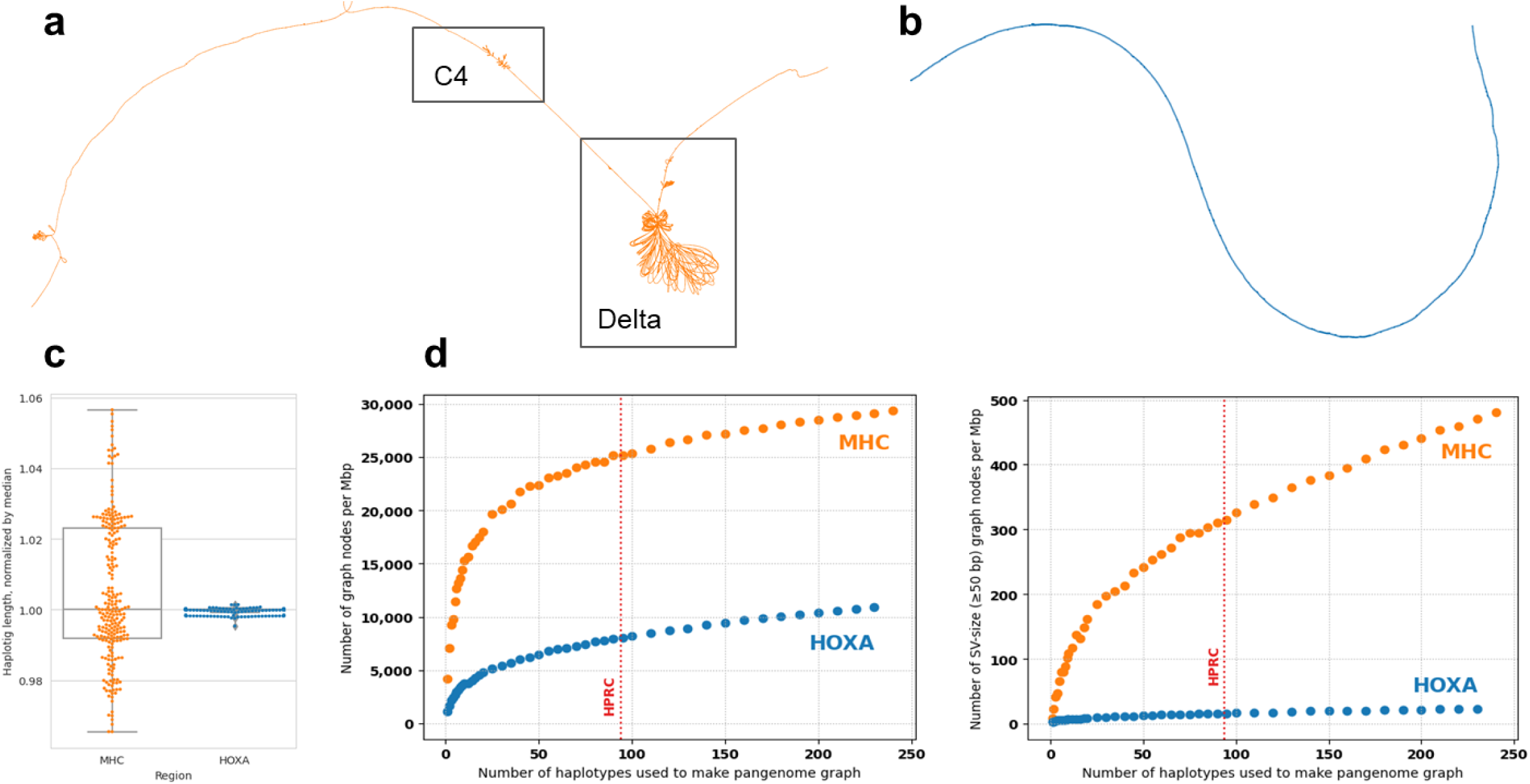
Building a pan-MHC graph. **a.** Bandage^66^ plot showing the pan-MHC graph. Clearly visible to the right is the delta polymorphic frozen block, which demonstrates significant structural diversity across assemblies. The C4 region is also noted, where there are several combinations of *C4A* and *C4B* long/short and CNVs. **b.** Bandage plot for the pan-HOXA graph, showing very little structural diversity relative to the similarly-sized MHC graph. **c.** Boxplot of haplotig lengths from the MHC (orange) and HOXA (blue) regions. Within each group, the haplotigs’ sequence lengths are normalized to the median length of the group. **d.** Pan-MHC graph size as a function of the number of haplotig sequences used to build it. Each point represents an average of ten pan-MHC graphs created from random subsets of the 240 (MHC) or 230 (HOXA) haplotigs. *Left*: Number of graph nodes per Mbp. *Right*: Number of graph nodes containing ≥50 bp of sequence per Mbp. The vertical dotted line represents the number of haplotypes released by the HPRC in Liao *et al*.^40^

**Table 4:** Superpopulation representation in our pan-MHC graph.

| Superpopulation | Number of haplotypes |
| --- | --- |
| African | 72 |
| American | 68 |
| East Asian | 44 |
| European | 12 |
| South Asian | 42 |
| Admixed | 2 |

The size and scope of our pan-MHC graph is heavily dependent on the number of haplotigs used to create it. To explore this dependence, we subsampled sets of haplotigs from our full set, built the subsets into pan-MHC graphs, and counted the nodes in these graphs. A graph’s total node count serves as a metric for its power to call small variants; similarly, its count of long-sequence nodes (≥50 bp) is a metric for calling structural variants. We found that, for the pan-MHC but not pan-HOXA graph, the node count continues to increase significantly with each added haplotig (**Fig. 5d**).

We used our reference graph creation method to perform an additional validation of the small variants in our assemblies. We generated a pan-MHC graph using only the two haplotigs of a contiguous diploid assembly for HG001 generated with HiFi reads. Then we deconstructed this graph to yield a set of phased variant calls, relative to the GRCh38 chromosome 6 reference. Comparing these assembly-based calls to the NIST v4.2.1 high confidence calls showed a sensitivity of 0.9922, precision of 0.9940, and F_1_ score of 0.9931.

### Our pan-MHC reference graph outperforms GRCh38 for variant calling

One of the most common applications of reference genomes is to align short-read datasets to them and call variants. Our pan-MHC graphs, in order to be usable as references, must be applicable for this task. To test this, we took a set of high-depth Illumina sequencing reads of HG001 and downsampled them to various depths from 1× to 95×. Then we aligned the downsampled readsets to our pan-MHC graphs using vg giraffe^56^, called variants using DeepVariant^48^, and compared the variant calls with high confidence HG001 variant callsets obtained from NIST (version 4.2.1). We found the best performance at 34× depth, with an average recall of 95.0%, precision of 99.2%, and F_1_ score of 97.1% (**Fig. 6**). For comparison, we performed the same analysis using a linear reference (GRCh38), aligning the same readsets with BWA-MEM^57^ and calling variants with DeepVariant. At all depth levels, the pan-MHC graph produced variant calls with equal or slightly higher F_1_ scores than the linear reference (**Fig. 6**). Thus our pan-MHC graph is not only serviceable for short-read alignment and variant calling, but actually more reliable than the state-of-the-art standard with a linear reference genome.

**Figure 6:**
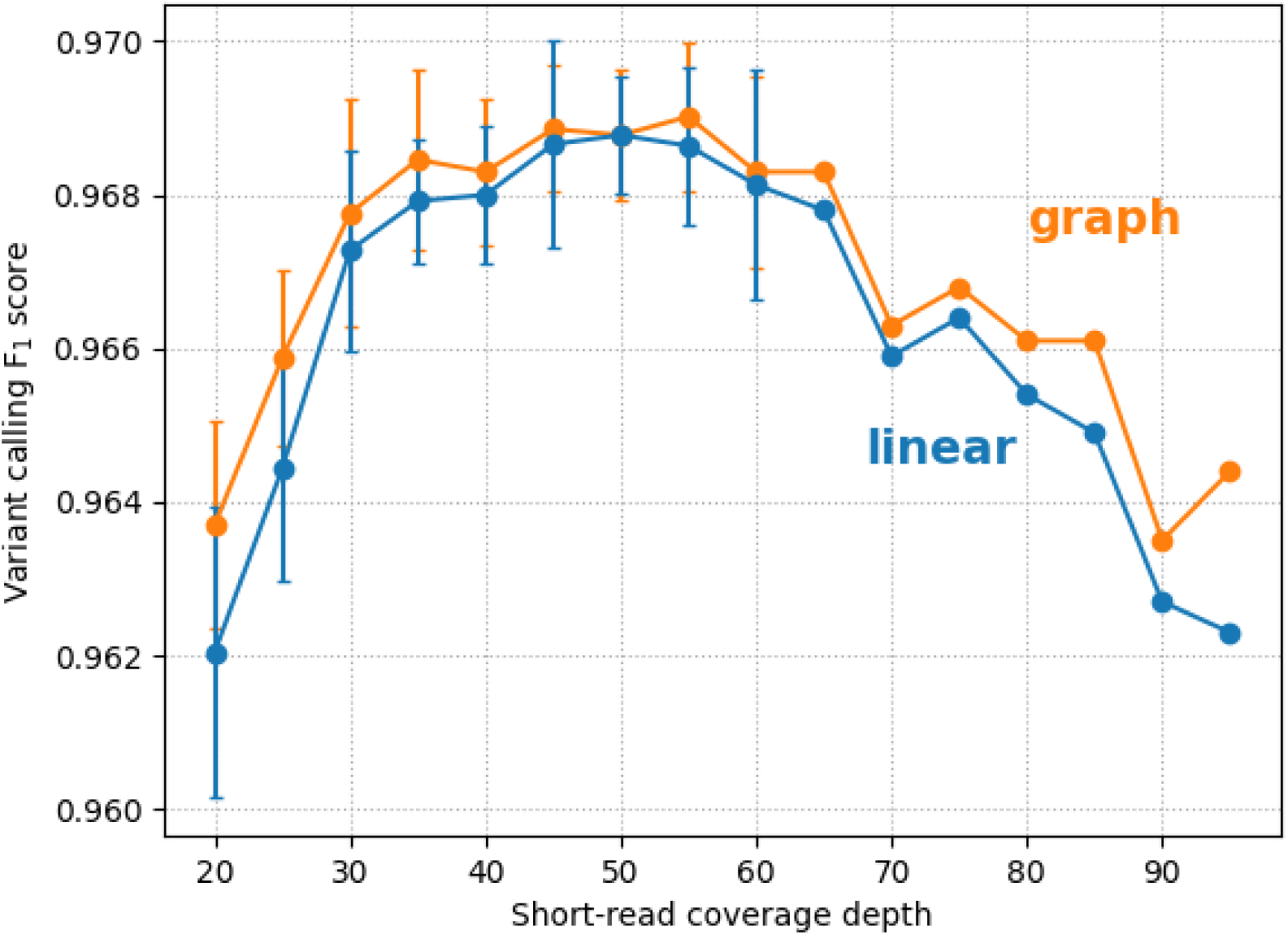
Pan-MHC graph outperforms linear reference genome for variant calling. Alignment of short reads to our pan-MHC graph (orange), in comparison with a standard approach using alignment to a linear reference genome with BWA-MEM^57^ (blue), produces improved variant calling *F1* scores across a range of input depths. *x*-axis: downsampled short-read coverage depth of HG001. *y*-axis: variant calling *F1* scores. Each data point represents a mean and standard deviation of 1-5 independent downsampling experiments.

## Discussion

Our collection of 246 highly accurate, fully contiguous, and phased full-length MHC sequences is, to our knowledge, unprecedented in both scale and population diversity. Our 1,246 putative novel allele sequences represent a potentially large step forward in our understanding of *HLA* sequence variation, especially at less well studied loci. Even among the 11 particularly well-studied *HLA* loci, we saw 271 putative novel allele sequences, of which 35 (12.9%) have coding sequence variants. Our pan-MHC reference graph based on 240 MHC haplotigs contains 10.3 Mb of sequence not represented in GRCh38 (**Fig. 5a**; **Table 1**) and represents more ancestral groups than previously reported thanks to the diversity of HPRC (**Table 4**). We should note that several of the novel alleles may have been observed in previous studies but not added to the IMGT/HLA database, and that the sequences are considered provisional pending orthogonal validation or review.

The pan-MHC graph is immediately applicable to genomic analyses in the MHC region, such as read alignment and variant calling. It can be used to impute MHC variants from existing short-read datasets, even WGS datasets with depth as low as 20× (**Fig. 6**). It already surpasses standard linear reference-based approaches, and we anticipate that it will continue to improve with ongoing advancements in the development of graph-based alignment methods^56,58^. Our graph has the potential to help downstream analyses, such as GWAS fine-mapping^10,13^ and biomarker discovery^11^.

Based on our comparative analysis, we find that our MHC assemblies of QBL and DBB are the most accurate to date (**Fig. 3b**; **Table 2**). For PGF we find that our assembly and the GRCh38 assembly are of comparable accuracy, and we find numerous putative errors in the human reference genome that has been in use for nearly a decade^59^. Finally, comparing our assemblies to those of HPRC we find a very high degree of sequence identity (**Fig. 3a**). One point of emphasis related to our assembly method is its utilization of only a single data type, PacBio HiFi sequence reads. In contrast to the HPRC’s use of trios for haplotype phasing^40^, we rely on the length of PacBio reads to bridge stretches of homozygosity. The prevalence of such stretches of homozygosity in MHC is a likely reason for phase loss in our assemblies relative to the HPRC’s. We anticipate that our efforts to increase our read lengths might mitigate this problem. Overall, our custom workflow for the MHC achieves excellent sequence contiguity and confident phasing over multi megabase regions using HiFi reads alone. Together with our targeting methods, this opens the possibility of routinely sequencing, assembling, and phasing the full MHC.

Our MHC haplotigs contain a large number of putative novel alleles and CNVs in the *C4A∼C4B* region (**Fig. 4a**; **Table 3**). These observations suggest a large degree of previously unreported allelic diversity and population substructure in the C4A∼C4B region. For instance, we show that some CNV of *C4A* and *C4B* are in LD with alleles of MHC genes, and that these haplotypes could be population-specific. In this way, our discoveries could help contextualize other research into C4 and add clinical value. For example, a previous study^18^ found that *C4A* and *C4B* CNVs correlate with schizophrenia risk; in light of our new results, it is feasible that the causal risk variants are in fact other alleles of the MHC region. To systematize our understanding of C4, it may be necessary to develop a more detailed nomenclature and terminology for C4 alleles and haplotypes; we invite the research community to discuss these considerations.

We expect the impact of the pan-MHC reference and graph to continue to grow with the number of available sequences and represented populations. Some of our analyses, such as our measurement of LD in C4 (**Fig. 4b**), were hamstrung by our relatively small sample size. Furthermore, even after this study, the diversity of the MHC region remains unrepresented in many human populations. Note that we could easily apply our method to assemble more sequence-contiguous MHC haplotigs, either by sequencing more cell lines or as more high-depth long-read sequencing datasets like HPRC’s become available. We have shown that the size of our MHC haplotig collection, significantly larger than preexisting collections, translates to greater capacity for variant calling (**Fig. 5d**), and this capacity would likely continue to grow if more MHC haplotypes were added. If the new haplotypes were to come from diverse populations, such as those being sequenced by the HPRC^40^, they would translate directly into an improved capacity for genomic analyses in these populations. This would benefit worldwide health equity, on account of the high clinical significance of MHC variation.

Lastly, we note that there are other human genomic regions beyond MHC that also contain clinically significant immune genes and are difficult to assemble, notably the killer-cell immunoglobulin-like receptor (KIR) region on chromosome 19^60–63^, the natural killer cell (NKC) region on chromosome 12^64^, and the immunoglobulin heavy chain (IGH) locus on chromosome 14^63^. While this study describes the work we performed around MHC, our platform is general and our target enrichment methods could readily be applied to any genomic region. We plan to target our method to other regions with clinical importance and sequence enough haplotigs of each region to create a true pan-immunogenome reference.

## Acknowledgements

The authors would like to thank Jill Hollenbach for very helpful discussions on the *C4A* and *C4B* genes, and Katsushi Tokunaga, Charles Khor Seik-Soon, and Jill Hollenbach for valuable discussions and collaboration on related work for the KIR region. Finally, we would like to thank Charles Lee, Qihui Zhu, and Feyza Yilmaz from The Jackson Laboratory for valuable feedback and discussions in an earlier MHC assembly collaboration, and Marcelo Fernández-Viña and Mary Carrington for encouragement and guidance when we embarked on this project.

## Conflict of interest disclosure

L.H., S.M.A., J.N.B., J.K.D., R.S.J., G.M.-S., H.M., B.D.T., and S.E. are employees and holders of shares or stock options in Base5 Genomics, Inc. A.J.M. is a shareholder in Base5 Genomics, Inc.

## Data availability

Our assemblies of the DBB, QBL, and PGF assemblies are publicly available in a Github repo (https://github.com/base5genomics/public-MHC-haplotigs). We are in the process of submitting these assemblies to NCBI Assembly.

## Supplementary Figures

**Supplementary Figure 1:**
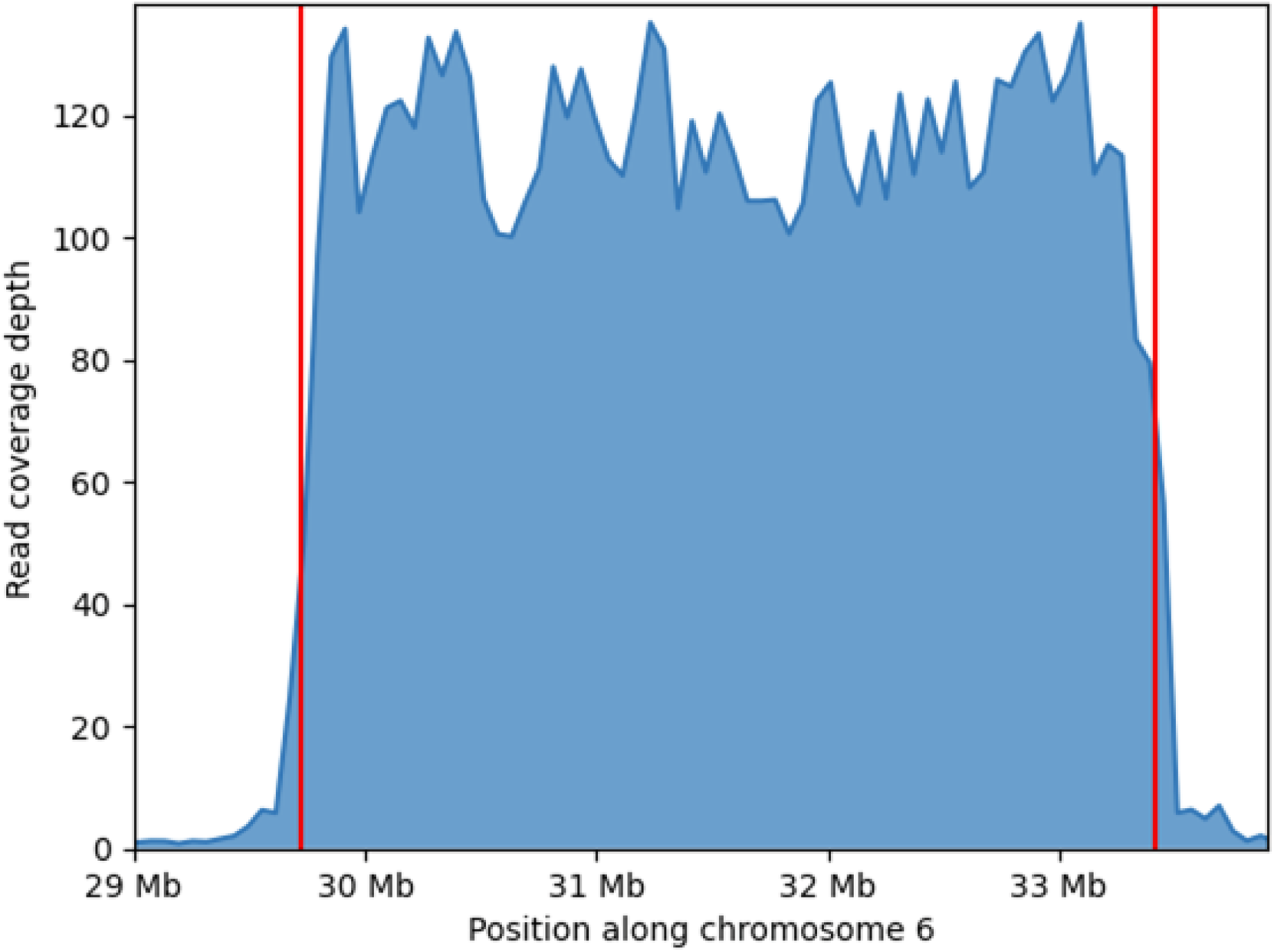
Enrichment of MHC region. Coverage depth plot of the MHC region of an MHC-enriched PacBio sequencing library of the PGF cell line, which achieved 93.7× enrichment (see **Methods**). Since this coverage plot is based on alignments of reads to GRCh38, and since MHC in GRCh38 is constructed from this individual (PGF), this depth plot provides an especially faithful representation of the depth, unaffected by reference bias.

**Supplementary Figure 2:**
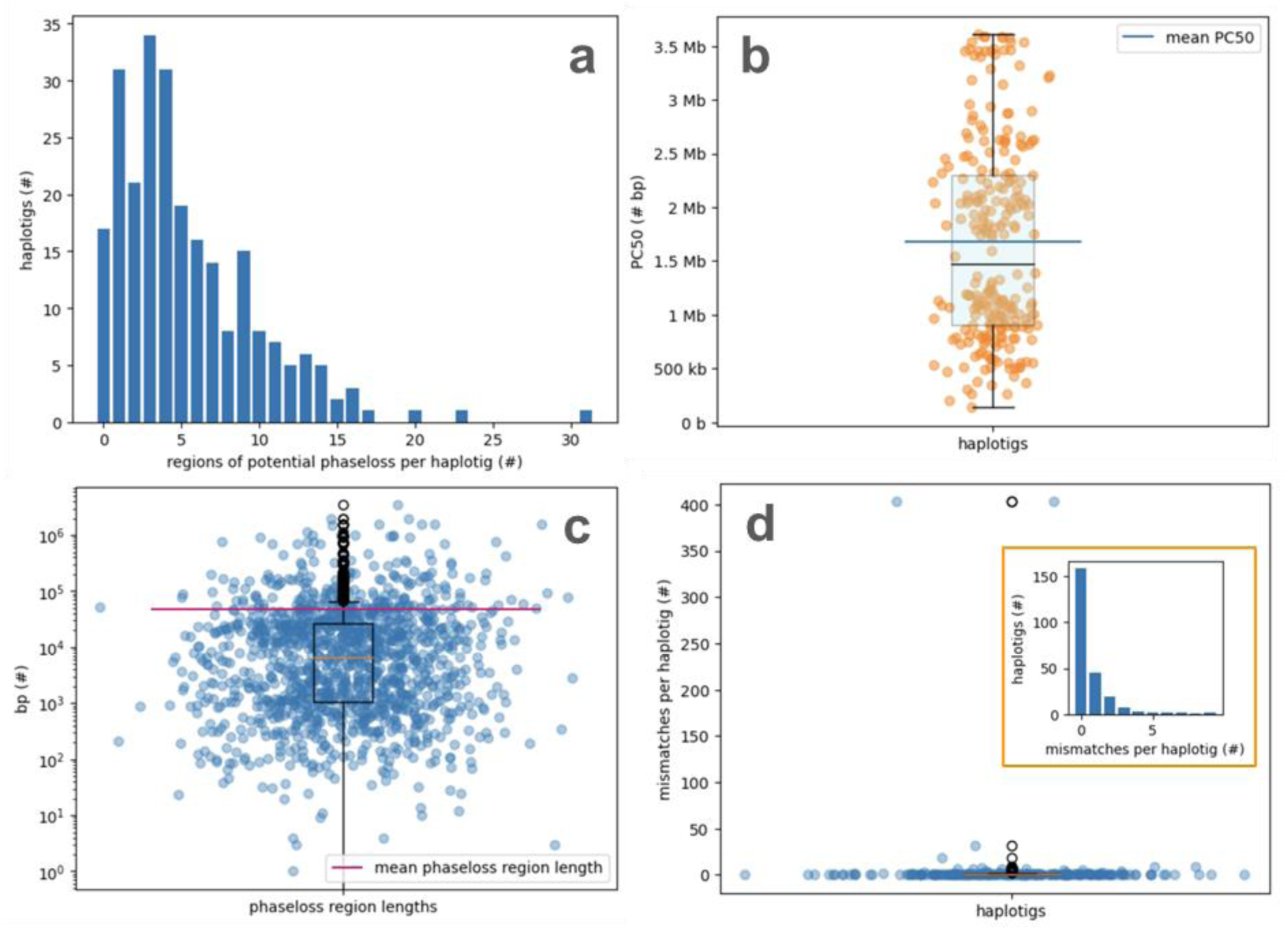
Inspection of our assembled haplotigs. **a.** Histogram showing the count of haplotigs (*y*-axis) having a given number of regions of potential phase loss per haplotig (*x*-axis), across 246 haplotigs. **b.** Box- and scatterplot showing PC50 values in bp. Each point represents one of our 246 assembled haplotigs; scatterplot represents exact PC50 values for each haplotig, with the 1st, 2nd, and 3rd quartiles shown in the boxplot. The horizontal blue line shows the mean value across haplotigs. **c.** Box- and scatterplot showing lengths of regions of potential phase loss in bp. Each point represents one of our 246 assembled haplotigs, with horizontal pink line showing the mean value across haplotigs. **d.** Box- and scatterplot showing the number of mismatches per haplotig (y-axis) having a given number of systematic mismatches (SMs, ≥51% of high-quality best-match aligning reads) across 246 haplotigs. Inset, orange-outlined histogram shows the distribution of the same data for only the 242 haplotigs with <10 SMs: count of haplotigs (*y*-axis) having a given number of mismatches per haplotig (*x*-axis).

**Supplementary Figure 3:**
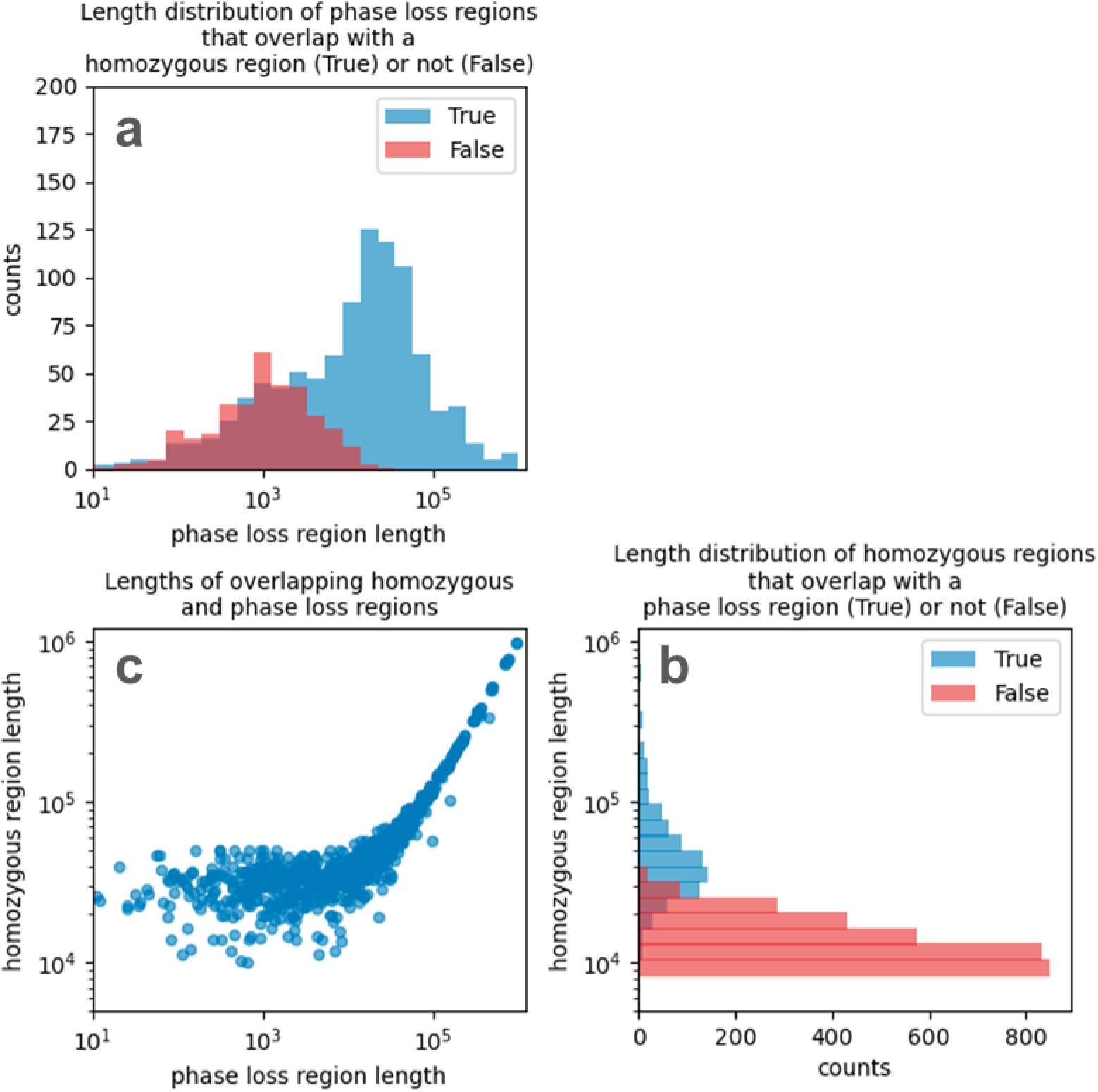
Phase loss correlates with homozygosity. **a.** Length distribution of phase loss regions, as identified by a drop in coverage depth of unambiguously aligning reads. Red: phase loss regions that do not overlap a homozygous region. Blue: phase loss regions that overlap a homozygous region. **b.** Histogram of the length distribution of homozygous regions, as identified by aligning the two haplotigs to each other. Colors are as in **a**. **c.** Scatterplot of the phase loss regions that overlap with a homozygous region. The respective lengths are plotted against each other.

**Supplementary Figure 4:**
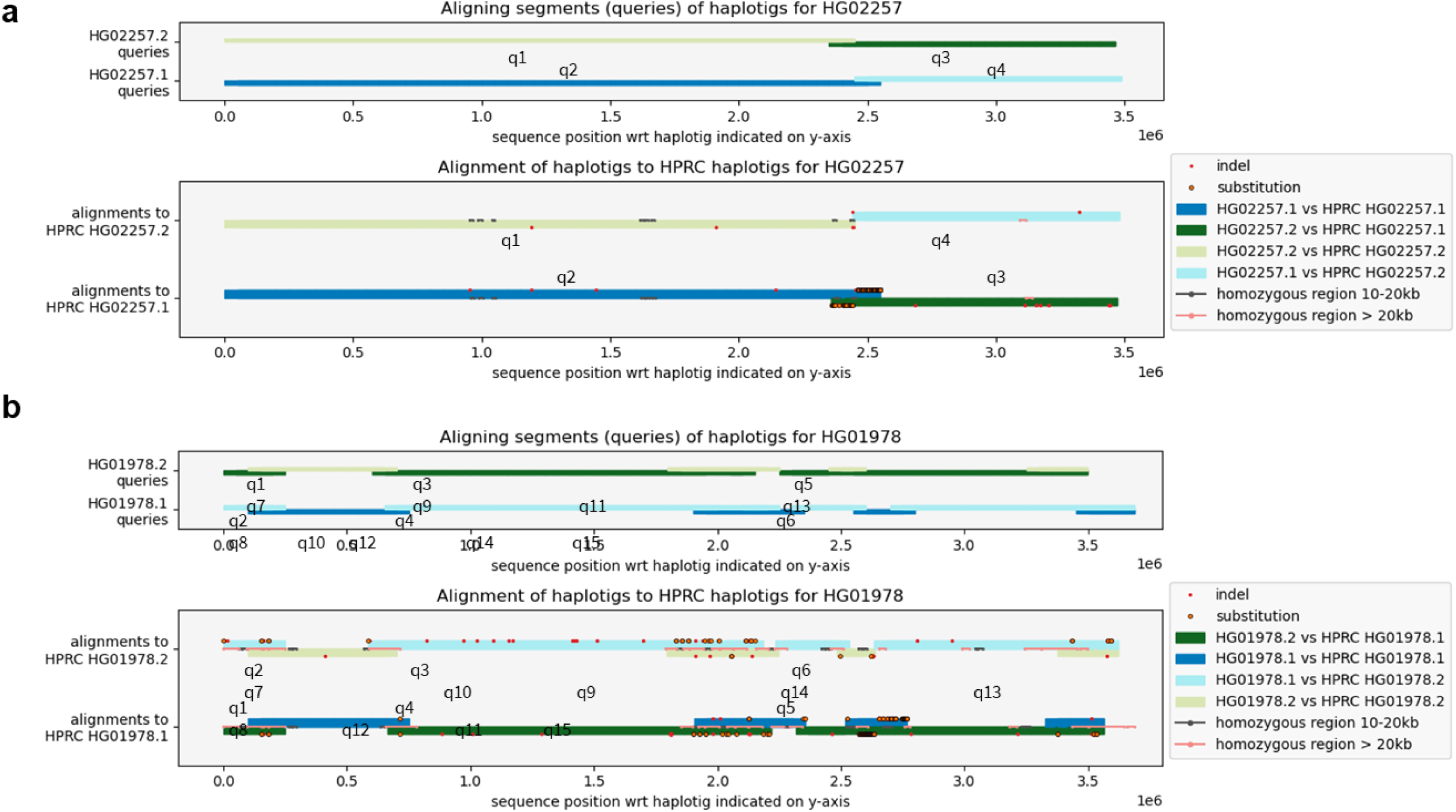
Phase switches in HPRC assembly comparison. Homozygous regions larger than the typical read length are prone to phase loss which could give rise to a phase switch. The figure shows the alignments of our haplotigs to HPRC’s MHC haplotigs for two samples. The top panel in each figure shows the query sequences (q1, q2, q3,…) taken from our haplotigs and the bottom track panel shows the alignments of these to HPRC’s haplotig 2 and 1 respectively. Indel and substitution mismatches are indicated as well as homozygous regions. **a.** The alignments reveal a phase switch around position 2.5Mb: here our haplotig 1 switches from resembling HPRC’s haplotig 1 better to resembling HPRC’s haplotig2 better and vice versa. The phase switch coincides with a short (<20kb) homozygous region. **b.** The alignments reveal as many as 7 phase switches, each of which coincides with a larger (>20kb) homozygous region.

**Supplementary Figure 5:**
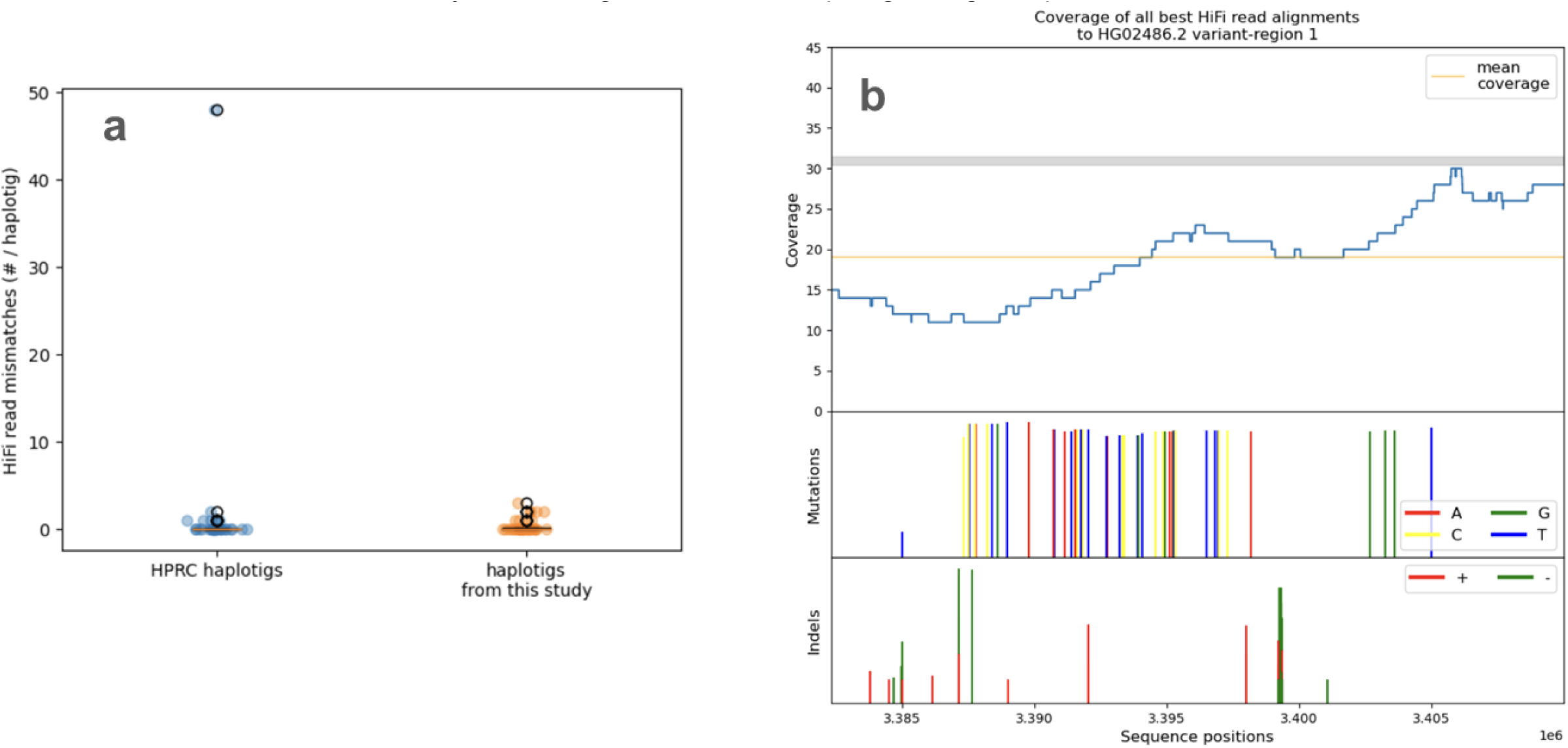
Assessment of our assemblies vs. HPRC assemblies. **a.** Box- and scatterplot showing number of mismatches per haplotig, with 36 HPRC assemblies in blue (left) and our 36 assemblies for the same 18 cell lines in orange (right). Each point represents one haplotig; scatterplot shows the number of mismatches for that haplotig. **b.** Coverage depth and mutations for the region of significant misassembly in HPRC-assemblied haplotig HG02486.2, which is the outlier point in subfigure **a**. The top panel shows coverage (blue line) and mean coverage (yellow line) of best-match high-quality HiFi read alignments to the haplotig; dropping coverage indicates that read alignment quality becomes very poor to the right end of the region. The middle and bottom panels show points at which HiFi reads have either a mismatch or indel, respectively, relative to the haplotig sequence; the height of a line at a given position is proportional to the relative number of disagreeing reads, and the full-height lines indicate that every read disagrees with the haplotig at a given position.

**Supplementary Figure 6:**
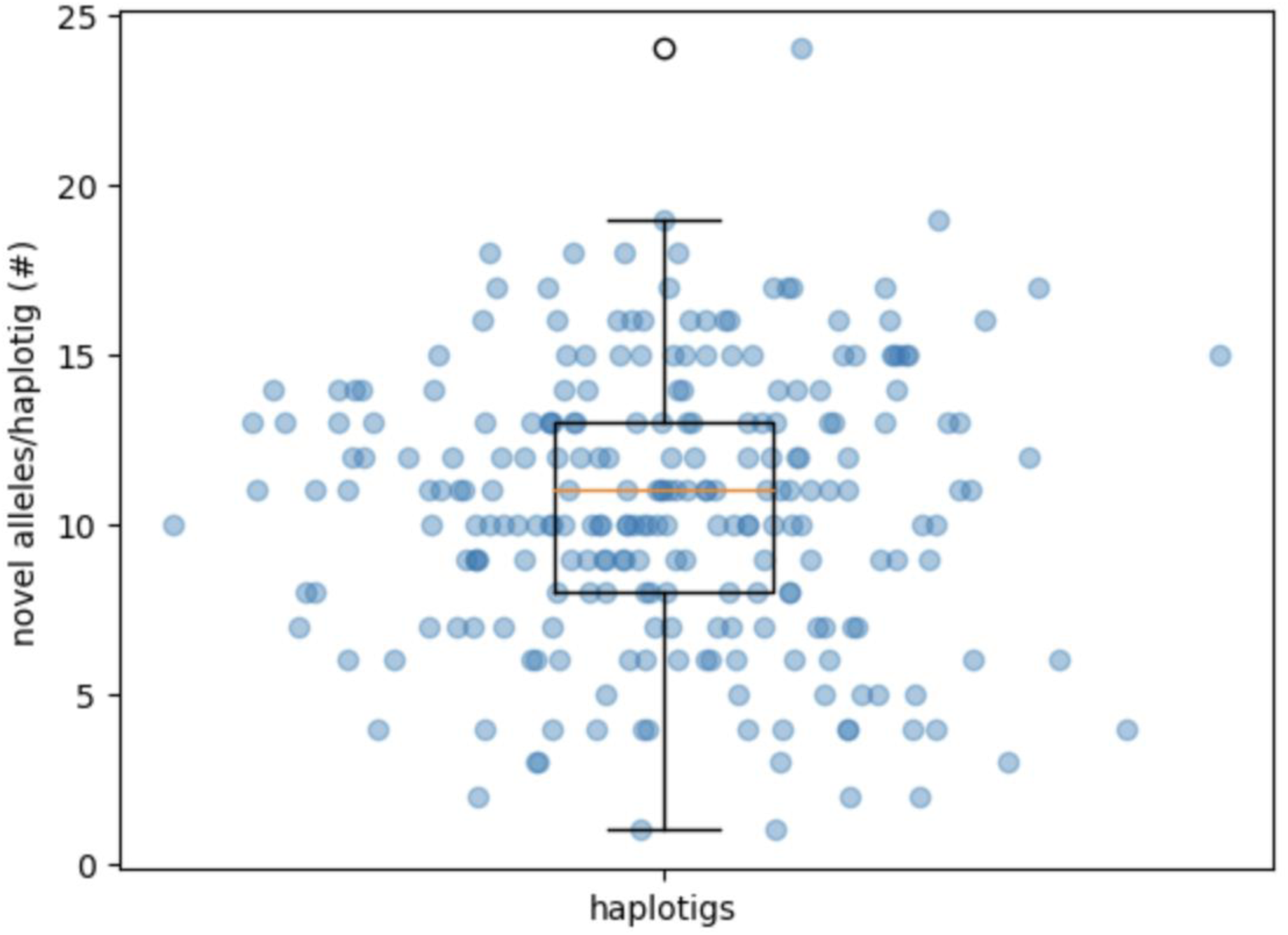
Novel allele distribution across haplotigs. Box- and scatterplot showing number of novel alleles called per haplotig. Each point represents one of our 246 assembled haplotigs; scatterplot shows the number of novel alleles called for that haplotig, with the 1st, 2nd, and 3rd quartiles shown in the boxplot.

**Supplementary Figure 7:**
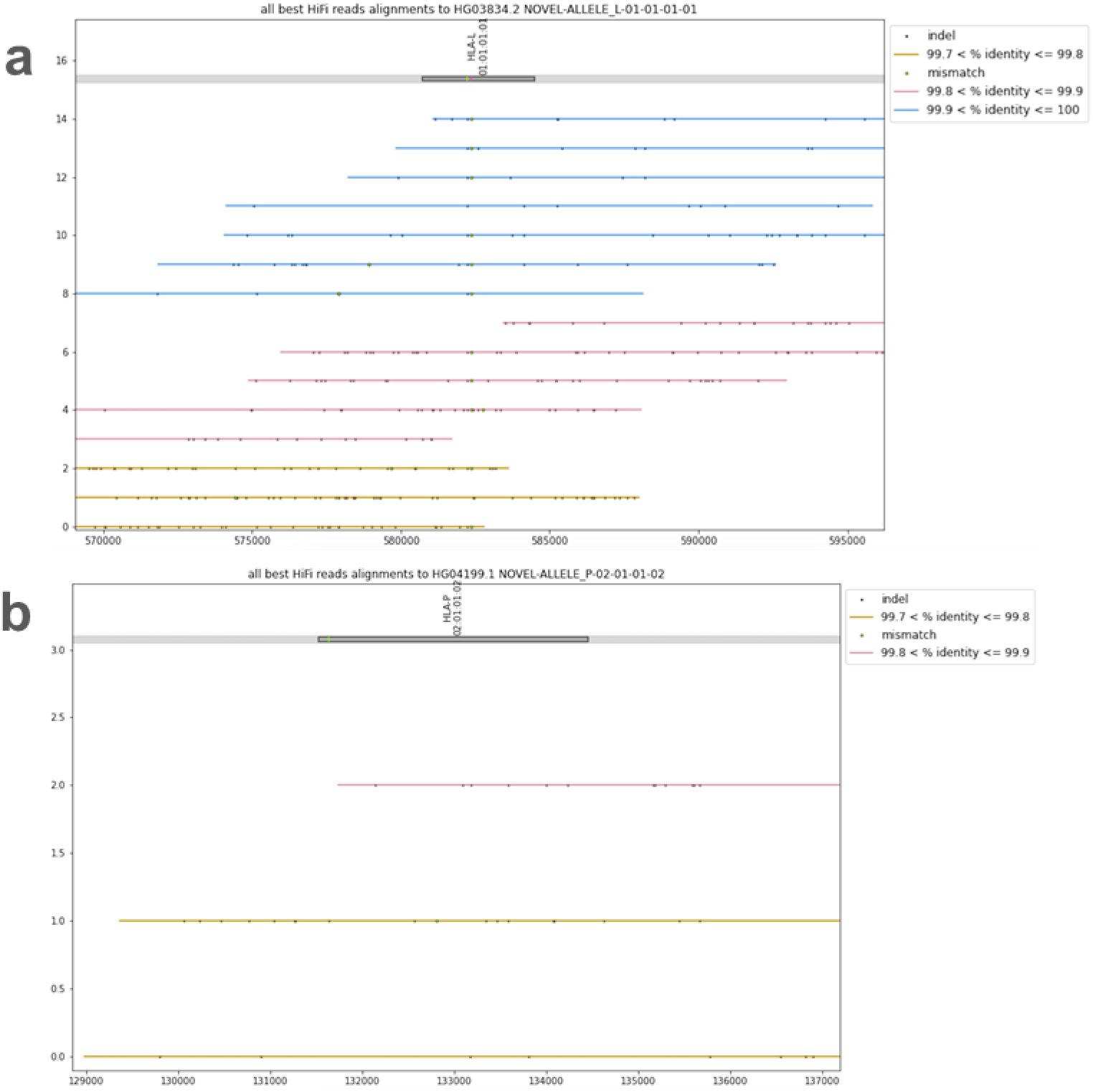
Examples of failed novel allele inspections. **a.** A putatively novel *HLA-L* allele for haplotig HG03834.2, has two variants, a mismatch and an indel, called against the database sequence for L*01:01:01:01 in the same positions where local point misassemblies are observed (around position 582,500). **b.** A putatively novel *HLA-P* allele for haplotig HG04199.1 has HiFi read depth that is too low for confidence in the allele call.

**Supplementary Figure 8:**
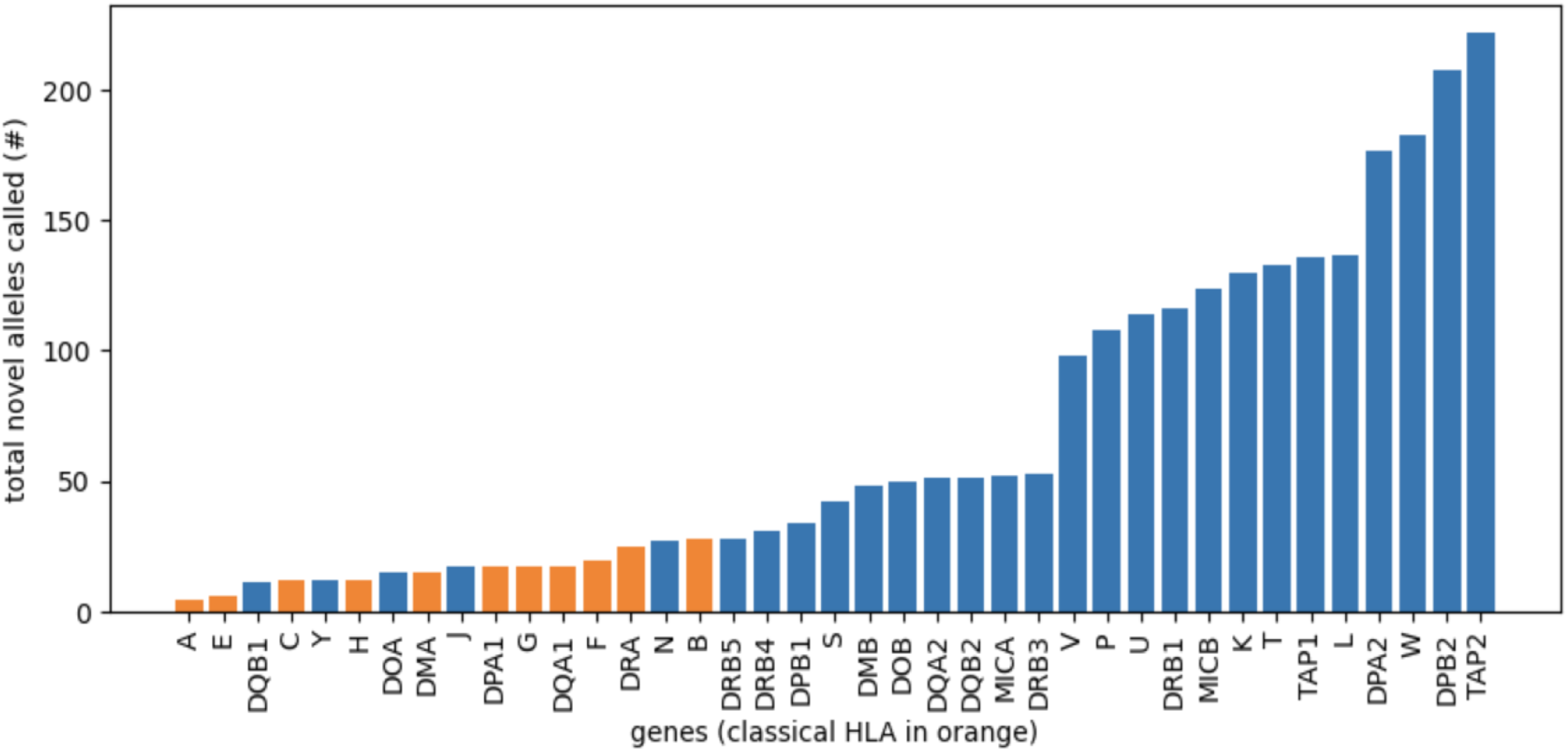
Frequency of novel alleles by locus. Barplot showing the number of times a variant was called against the IMGT/HLA allele database^17^, separated by locus. The classical HLA loci, which are relatively well characterized and are typically captured in HLA typing kits^26^, are shown in orange.

**Supplementary Figure 9:**
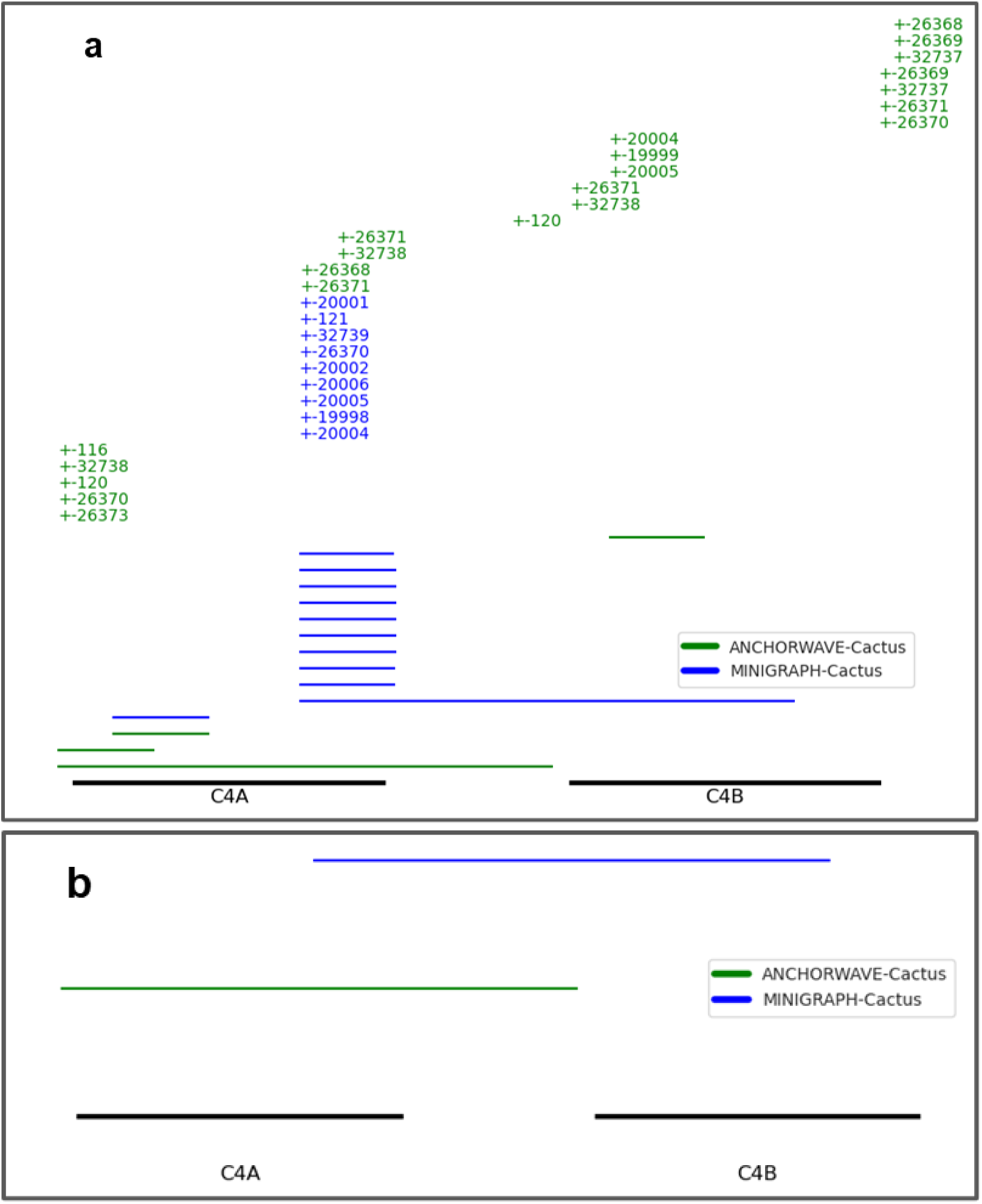
Graph deconstruction in C4 region in AnchorWave-Cactus vs. Minigraph-Cactus. **a.** Structural variant calls from the deconstructed Minigraph-Cactus graph (blue) vs. the AnchorWave-Cactus graph. Deletions relative to reference are shown as continuous horizontal lines. Insertion start positions are shown with a + along with the length of the insertion. The Minigraph-Cactus graph has no clear full *C4A* or *C4B* deletions and multiple unexpected deletions and insertions, while the AnchorWave-Cactus graph properly shows a whole-gene deletion and ‘short’ *C4A* and *C4B* genes. **b.** Comparison with a single sample (HG00733) in the same region shows the expected C4 copy-count in the AnchorWave-Cactus graph vs. an unexpected *C4A*+*C4B* hybrid in the Minigraph-Cactus graph.

## Methods

In all analyses involving the human reference genome, we used version GRCh38.p12.

### Library preparation and sequencing

We purchased the cell lines DBB, QBL, PGF, and HG00733 from Coriell Institute or Sigma Aldrich and cultured them at the Cell Culture Facility at UC Berkeley. We embedded the cells in agarose plugs and subjected them to Base5 Genomics’ proprietary enrichment protocol, including Cas9-based methods, to target the MHC region. We used the enriched high molecular weight DNA to prepare SMRTBell libraries and sequenced them and whole-genome libraries using Pacific Biosciences’ Sequel II instrument, creating HiFi sequence reads.

### Pre-processing pipeline

To calculate metrics like coverage and enrichment, and prepare reads for our assembly pipeline, we first ran those reads through a read pre-processing pipeline. We aligned reads with minimap2^65^ version 2.2x (with parameters `-N 10 -p 0.1 --mask-level=0.5 -O5 -E2 -A1 -B3`) to GRCh38. These alignments were used as the input to the assembly pipeline. To continue on with the read processing pipeline, we extracted from these alignments all reads that aligned to the MHC portion of chromosome 6 or any of the MHC alternates. We then realigned those reads (using minimap2^65^ with the same parameters as before) to a reference that was GRCh38 but with all alternates removed. We then ran `samtools depth` (parameters `-a -r`) on those alignments, choosing as a region the MHC portion of chromosome 6. Importantly, the `samtools depth` command only considers primary alignments, thus preventing it from overestimating coverage in the region. From the result of that depth calculation, we obtained and plotted coverage in the MHC, averaging the coverage in 60kb bins to improve the plots’ legibility. We calculated enrichment as the coverage in the targeted region divided by the average coverage across the entire genome. We estimated average coverage across the entire genome as total read length divided by total genome length.

### Sourcing publicly available read datasets for creating assemblies

We selected publicly available PacBio WGS datasets from NCBI Sequence Read Archive (https://www.ncbi.nlm.nih.gov/sra/). We applied the following search filters: Organism = Homo sapiens; Access = Public; Platform = PacBio SMRT; instrument = Sequel II; Strategy = Genome. We further filtered the search result as follows: Total Bases ≥ 90,000,000,000; Submitter = “UCSC GI” or “University of California, Santa Cruz”. For each of the samples passing these filters, we downloaded a set of reads for this sample in `fastq.gz` format. Some datasets could not be downloaded due to unspecified server problems. We ultimately downloaded reads for a total of 131 samples.

### Diploid assembly overview

We developed a targeted diploid assembly workflow (**Fig. 1**), which produces highly contiguous and often fully phase-resolved targeted assemblies. This workflow incorporates multiple assembly steps that iteratively separate reads into groups corresponding to their respective haplotypes. We also developed a method of information injection, where reference sequences (*e.g.*, GRCh38 chr6, ALT contigs, etc.) and a primary haploid assembly provide the initial scaffold that allows a preliminary diploid assembly to be scaffolded based on phase. Briefly, we generate a primary haploid assembly with Canu^67^, from which we call variants with PEPPER-DeepVariant^68^, phase them with WhatsHap^69^, and then use this assembly to tag reads for secondary assembly with Canu^67^ and hifiasm^70^.

### Allele calling

We called alleles at a total of 39 loci, including 35 HLA loci (*HLA-A*, *HLA-B*, *HLA-C*, *HLA-E*, *HLA-F*, *HLA-G*, *HLA-H*, *HLA-J*, *HLA-K*, *HLA-L*, *HLA-N*, *HLA-P*, *HLA-S*, *HLA-T*, *HLA-U*, *HLA-V*, *HLA-W*, *HLA-Y*, *HLA-DMA*, *HLA-DMB*, *HLA-DOA*, *HLA-DOB*, *HLA-DPA1*, *HLA-DPA2*, *HLA-DPB1*, *HLA-DPB2*, *HLA-DQA1*, *HLA-DQA2*, *HLA-DQB1*, *HLA-DQB2*, *HLA-DRA*, *HLA-DRB1*, *HLA-DRB3*, *HLA-DRB4*, *HLA-DRB5*) and 4 non-HLA loci (*MICA*, *MICB*, *TAP1*, *TAP2*). This list was based on filtering the IMGT/HLA database^17^ for non-partial alleles with full gene sequences available. We called alleles in the *C4A* and *C4B* region using a separate process (see below).

Contiguous assemblies allow spatial annotation of allele calls, *i.e.*, exact delineation of the relative positions of genes and alleles. The problem of spatial allele calling is relatively straightforward, but it is not trivial. Genes in the MHC region have considerable sequence homology with one another and among their alleles, confounding the identification of sequences. Furthermore, we found that many assemblies contained genes without a perfect allele match in the database, necessitating a method to select a ‘best fit’ allele along with variations relative to that best match. To address these challenges, we developed a novel method for annotating diploid assemblies with high-confidence allele calls and precise assembly-oriented boundaries. We first aligned allele sequences sourced from the IMGT/HLA database^17^ version 3.53 to the diploid assembly using minimap2^65^. Next, we consider each output alignment which provides a candidate gene, allele, query length, query start, query end, reference start, reference end, and several parameters encoding alignment quality and the long-form cs string. We also calculated identity *I* as the quotient of fields 10 (number of matches) and 11 (alignment length) and coverage depth *C* as the quotient of query alignment length and the total query length. In this context, the reference is the diploid assembly haplotig and the query is a specific allele corresponding to a gene in MHC (*e.g.*, *HLA-B*, *HLA-A*, *MICA*).

We determined “best fit” allele calls by solving the following Integer Linear Programming (ILP) problem, which seeks to maximize the total weighted identity and coverage of all alleles subject to some basic constraints:

Maximize: 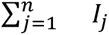

where *j* = 1…*n* are the alignments and *I_j_* are their sequence identities subject to:

1. Genes may not be selected more than a set copy number.
2. Allele positions must be mutually exclusive, such that their alignments do not overlap

We output allele calls into VCF files, then post-processed them to provide the arrangement of allele calls and boundaries on each haplotig in the assembly. We also annotated allele calls with small variant calls relative to the allele sequence, which are used to denote novel allele candidates.

### Inspection and validation of assemblies

We ran a number of experiments to assess the quality of our diploid assembly sequences and our novel allele calls. For each of the 123 samples, we aligned the MHC-region selected reads to the sample’s two assembled haplotigs with minimap2^65^ using parameters `-x map-hifi --cs`. We filtered alignments for length >500 bp and % identity with the assembled sequence >99%, where % identity is the percent of identical positions between the read and the assembly to which it is aligned, divided by the length of the alignment. For each remaining read, we chose one or more “best” alignments by first selecting those where the length was ≥95% of the read’s longest alignment or the alignment was not a partial alignment only, and then accepting all alignments with the lowest # of sequence mismatches against the assembly. We labeled reads with multiple equally good alignments as ambiguously-aligning, and we labeled those with a single best alignment unambiguously-aligning.

We then performed the following 5 assessments for each haplotig individually, as well as for its 5 polymorphic frozen blocks (PFBs) and for all alleles called with variants against the IMGT/HLA database sequence. First, we examined reads with point mismatches against the assembled sequence. We marked haplotig sequence positions where at least 51% of reads had a different base as having a potentially incorrect base, termed systematic mismatches (SMs). Secondly, we tested whether these potential mismatch positions clustered together in a region of high mismatch density, where a cluster of incorrectly called bases is indicative of misassembly. Thirdly, we checked overall read depth across the length of the haplotig or haplotig sub-region; we flagged areas with depth dipping below either 30% of mean region depth or 3 reads, whichever was larger. We additionally flagged assembled haplotigs or haplotig sub-regions with more than 5% of the region length falling under the low depth threshold defined in the previous sentence. Finally, we checked for regions of potential phase loss by looking at the unambiguous read alignments to each haplotig. In regions of homozygosity, reads cannot align unambiguously and so there are local dips in unambiguously aligning read depth; we flagged these for potential phase loss. As another method to identify potential phase loss, we identified homozygous regions directly by aligning the two haplotigs in an assembly to each other and finding >10kb (kilobase) identical regions. To guarantee that all homozygous regions are found we used alignments of a 100kb sliding window along one haplotig to the full length sequence of the other haplotig.

### Definition of PC50

We define a new metric, *phase contiguity 50* (PC50), which we base on the sequence contiguity metric N50. PC50 is defined as the length *X* at which at least half the haplotigs’ bases are in phase contiguity with ≥*X* bases. To calculate *X*, we first identify phase-contiguous segments within each fully sequence-contiguous haplotig. These fall into two categories: 1) each “phased region” across which there are over 3 high-quality, unambiguously-located best-match read alignments; and 2) the “potential phase-loss regions,” with less than 3 unambiguous best-match read alignments that link the phased regions. In this way, the entire haplotig is segmented into alternating phased heterozygous and potentially homozygous regions, each of which is confidently phase-contiguous within its boundaries. *X* is found for a given haplotig by determining the smallest subset of phase-contiguous segments that cover half of the haplotig length, then setting *X* to be the length of the shortest segment in this subset. Note that each heterozygous region is fully in phase with its neighboring homozygous regions, and separating these regions generates conservatively low phase-contiguous region lengths.

### External assemblies used in validation

We used the following publicly available existing MHC assemblies for comparison:

- PGF: The MHC region extracted from chr6 of GRCh38^47^.
- QBL and DBB: Houwaart *et al*.^32^
- HPRC samples: The MHC region, defined as GRCh38 chr6 position 28510120-33480577 was found to be contiguous in 25 out of 46 diploid HPRC^40^ samples. We selected 18 of those samples to compare with our assemblies in this publication.

For all samples we restricted the comparison to the ∼3.5Mb region, defined by chr6:29,668,442-33,184,109 on GRCh38, that falls within our targeted region (**Supp. Fig. 1**). For PGF, QBL, and DBB, the external assemblies have only one haplotig and comparison is done by alignment of our haplotigs to the external haplotigs with minimap2^65^. For the HPRC samples where both our assemblies as well as the external assemblies are diploid the comparison is complicated by occasional phase switches. We addressed this by first identifying phase switches using alignments of a 100kb sliding window along our haplotigs to find regions where the sequence of our haplotig 1 switches from resembling HPRC’s haplotig 1 better to resembling HPRC’s haplotig 2 better and vice versa. Further sequence identity comparison was then done by locally excluding the phase switch regions. This reduced the comparable sequence to 96% of the haplotig length on average. Finally, we used the self-alignment method described earlier to identify larger homozygous regions (>15kb).

### DBB/QBL/PGF comparison vs. external assemblies

We aligned to the assemblies using the following readsets from SRA:

- DBB: <u>SRX19214161</u> (PacBio), <u>ERX525931</u> (Illumina)
- QBL: <u>SRX19214164</u> (PacBio), <u>ERX525979</u> (Illumina)
- PGF: <u>ERX525976</u> (Illumina)

We chose to use only one haplotig from each of our diploid MHC assemblies in order to make the comparison even (since the external MHC assemblies only have one haplotig), and also because the two haplotigs each of our diploid MHC assemblies are much more similar to each other than either is to the haplotig from the external MHC assembly. We also clipped the external MHC assemblies to ensure that they span the exact same portion of chr6 as our MHC assemblies. For each cell line, we constructed a full reference genome by patching the assembly into chr6 of GRCh38 and excluding alt contigs. We aligned short reads to these references with BWA-MEM^57^ (version 0.7.17), then used these alignments to call variants with DeepVariant^48^.

### *C4A* and *C4B* allele calling and nomenclature

*C4A* and *C4B* gene sequences differ in three key ways: (1) whether they are isotype *A* or *B*; (2) whether they code for proteins that produce blood group Rodgers (*Rg)* or Chido (*Ch*); and (3) whether they are long form or short form. *A/B* and *Rg/Ch* are distinguished based on sets of SNPs while long/short are distinguished based on the length of the gene, as follows:

1. There is a 17bp region on *C4A/C4B* which, by definition, has seq-A ‘CCTGTCCAGTGTTAGAC’ on *C4A* and has seq-B ‘TCTCTCCAGTGATACAT’ on *C4B*. There are 5 SNP differences between these sequences. This region begins at position 7667 if the gene is short form, or position 14027 if the gene is long form. We call an allele *A* if it contains seq-A exactly once and does not contain seq-B. Conversely, we call an allele *B* if it contains the seq-B exactly once and does not contain seq-A.
2. Occurring 456 bp downstream, at position 8123 if short form or 14483 if long form, is a 16bp region which, by definition, has seq-Rg ‘GCCTGTGGACCTGCTC’ for blood group *Rg* and seq-Ch ‘CCCTGCGGACCTGCGG’ for blood group *Ch*. There are 4 SNP differences between these sequences. We call alleles as *Rg* or *Ch* using the same logic as above.
3. *C4* alleles come in long form (21kb) or short form (14.6kb). The long form’s extra length comes from an endogenous retroviral sequence that has integrated into intron 9^71^. We determined long versus short at the same time we initially located these alleles on the haplotigs, by using minimap2^65^ to align both long and short forms of the gene to the haplotig. Consistently, either the long form had >99% identity and the short form had <70% identity, or vice versa; therefore this method uniquely located and identified the form of every *C4* allele.

As *A* typically is associated with blood group *Rg*, we denote an allele that has both as simply *C4A*. As *B* typically is associated with blood group *Ch*, we denote an allele that has both as simply *C4B*. We denote *A* co-occurring with *Ch* as *C4recA-Ch* where “rec” stands for “recombinant”. Similarly we denote *B* co-occurring with *Rg* as *C4recB-Rg*. To indicate long or short isoform, we append “-L” or “-S” to the name, for example, *C4A-*L.

There were some rare cases in which the above rules could not be cleanly applied. In these cases we applied the following nomenclature:

- Twice we observe an allele that is clearly “L” and clearly *Rg*, but is not an exact match to either *A* or *B*. It has a sequence ‘GCTGTCCAGTGTTAGAC’. Since this sequence differs from seq-A by only one base (the first base, which on seq-A is ‘C’), a base which does not match seq-B either (which has ‘T’ as a first base), we call this allele as an alternate of *A*, in this study referred to as *C4A*-L alt1.
- Once, we observe an allele that is clearly *A* and clearly “L”, but is not an exact match to either *Rg* or *Ch*. It has a sequence ‘GCCTGTGGACCTGCGG’. We denote this allele C4*A-*?-L alt2.
- Once, we observe an allele that is clearly *A* and clearly “L”, but is not an exact match to either *Rg* or *Ch*, in a different way from the previously described allele. It has a sequence ‘GCCTGCGGACCTGCTC’. We denote this allele C4*A-*?-L alt3.

To address the possibility that HiFi reads might not be long enough to fully resolve tandem duplicate copies of *C4A∼C4B* that exceed two in a row, we used depth of coverage along the *C4A∼C4B* region to infer additional copies of the same gene.

### Linkage disequilibrium (LD) between *C4A∼C4B* and other genes within the MHC

We tested the LD between the CNVs of the *C4A∼C4B* region and the 39 MHC loci. We used Arlequin^72^ (version 3.5.2.2) to compute *D’* and *r^2^* coefficients, and to test the significance of the exact test of LD between alleles utilizing a Markov chain of 1,000,000 steps and a dememorization process of 1,000,000 steps.

### Pan-MHC graph creation

We modeled our graph creation approach on that of the HPRC^73^. The HPRC’s bioinformatic method, called Minigraph-Cactus workflow, includes a graph creation process that is extremely robust and computationally efficient. However, we found that it produced suboptimal alignments in complex regions within the MHC. This was especially pronounced in the C4 region, where CNVs are extremely common, clinically relevant, and difficult to properly align to the reference which only contains one copy of the long isoform of each gene. Challenging alignments are also present in other highly variable regions of the MHC, which might hinder proper SV calling in downstream applications. Therefore, we altered the Minigraph-Cactus workflow, replacing the minigraph SV graph generation and alignment step with end-to-end Anchorwave^74^ (version 1.2.1) alignments with custom adaptation to properly connect the alignment outputs to `cactus_consolidate`.

Similar to Minigraph-Cactus, we generate an output ‘full graph’ containing all nodes and edges from every component sample. We also generate an analysis set, which clips nodes >10kb that are not part of the reference and nodes that are not supported by at least 2 component haplotigs (d2 analysis set).

### Graph-based short read alignment and variant calling

We aligned short reads from Illumina paired-end sequencing to the graph with Giraffe (vg 1.49.0)^56^. We generated Giraffe indices from the d2 analysis set with a maximum node size limited to 100 bp, which we found slightly improves alignment and variant calling relative to the default node size of 1024 bp.

We surjected alignments from graph space to linear space (GRCh38 chr6 reference) with vg^56^. Next, we left aligned the bam alignments with bamleftalign (freebayes^75^ version 1.3.6) and realigned indels with abra2^76^ (version 2.24). We trained a custom DeepVariant^48^ (version 1.5.0) model using truth calls from the NIST (National Institute of Standards and Technology) Genome in a Bottle (GIAB) release 4.2.1 for HG006 based on alignments to our graph genome, then called small variants using our custom model. We normalized small variants with bcftools^77^ (version 1.17).

### Public datasets for short-read variant calling

For training models and validation of short-read variant calls, we obtained high-confidence variant calls, short reads, and PacBio HiFi reads from the NIST GIAB samples NA12878 (HG001) and NA24694 (HG006). We used version 4.2.1 of the high-confidence variant calls for training and evaluation of our variant calling methods.

112 of our 120 samples (**Supp. Table 1**) were cell lines from the 1000 Genomes Project. For these 112 samples, we obtained paired-end short reads for the MHC region from the 30× depth WGS reads generated by Byrska-Bishop *et al.^53^*. We extracted short reads from the aligned CRAM files that mapped to the MHC region on chr6, HLA allele sequences, or MHC alt haplotypes.

Data reuse statement for 1000 Genomes: https://www.internationalgenome.org/data-portal/data-collection/30x-grch38

### Linear reference-based short read alignment and variant calling

We aligned short reads to the GRCh38 chr6 reference, excluding alts, with BWA-MEM^57^. We called variants with DeepVariant^48^ using the WGS built-in model.

### Control region

We defined a control genomic region around the *HOXA* gene cluster: chr7:25296538-29630479. We find that this region exhibits high conservation across primates (as measured by the “30 mammals conservation by PhastCons (27 primates)” track on the UCSC GenomeBrowser which can be added by clicking “Cons 30 Primates” under the “Comparative Genomics” heading), exhibits low repeats (as measured by the “RepeatMasker” track on the UCSC Genome Browser), and has a GC content similar to MHC.

## Notes

https://github.com/base5genomics/public-MHC-haplotigs

## References

1. Chin, C.-S. et al. A diploid assembly-based benchmark for variants in the major histocompatibility complex. Nat. Commun. 11, 4794 (2020).

2. Horton, R. et al. Gene map of the extended human MHC. Nat. Rev. Genet. 5, 889–899 (2004).

3. Trowsdale, J., Ragoussis, J. & Campbell, R. D. Map of the human MHC. Immunol. Today 12, 443–446 (1991).

4. Campbell, R. D. & Trowsdale, J. Map of the human MHC. Immunol. Today 14, 349–352 (1993).

5. Krensky, A. M. The HLA system, antigen processing and presentation. Kidney Int. Suppl. 58, S2–7 (1997).

6. Cruz-Tapias, P., Castiblanco, J. & Anaya, J.-M. Major histocompatibility complex: Antigen processing and presentation. (El Rosario University Press, 2013).

7. Thorsby, E. A short history of HLA. Tissue Antigens 74, 101–116 (2009).

8. Moyer, A. M. & Gandhi, M. J. Human Leukocyte Antigen (HLA) Testing in Pharmacogenomics. in Pharmacogenomics in Drug Discovery and Development (ed. Yan, Q.) 21–45 (Springer US, 2022). doi:10.1007/978-1-0716-2573-6_2.

9. Johnson, A. D. & O’Donnell, C. J. An open access database of genome-wide association results. BMC Med. Genet. 10, 6 (2009).

10. Kennedy, A. E., Ozbek, U. & Dorak, M. T. What has GWAS done for HLA and disease associations? Int. J. Immunogenet. 44, 195–211 (2017).

11. Augusto, D. G. et al. A common allele of HLA is associated with asymptomatic SARS-CoV-2 infection. Nature (2023) doi:10.1038/s41586-023-06331-x.

12. de Bakker, P. I. W. et al. A high-resolution HLA and SNP haplotype map for disease association studies in the extended human MHC. Nat. Genet. 38, 1166–1172 (2006).

13. Naito, T. et al. A deep learning method for HLA imputation and trans-ethnic MHC fine-mapping of type 1 diabetes. Nat. Commun. 12, 1639 (2021).

14. Turner, T. R. et al. Widespread non-coding polymorphism in HLA class II genes of International HLA and Immunogenetics Workshop cell lines. HLA 99, 328–356 (2022).

15. Gaudieri, S., Leelayuwat, C., Tay, G. K., Townend, D. C. & Dawkins, R. L. The major histocompatability complex (MHC) contains conserved polymorphic genomic sequences that are shuffled by recombination to form ethnic-specific haplotypes. J. Mol. Evol. 45, 17– 23 (1997).

16. Norman, P. J. et al. Sequences of 95 human haplotypes reveal extreme coding variation in genes other than highly polymorphic and. Genome Res. 27, 813–823 (2017).

17. Barker, D. J. et al. The IPD-IMGT/HLA Database. Nucleic Acids Res. 51, D1053–D1060 (2023).

18. Sekar, A. et al. Schizophrenia risk from complex variation of complement component 4. Nature 530, 177–183 (2016).

19. Zhou, D. et al. Human Complement C4B Allotypes and Deficiencies in Selected Cases With Autoimmune Diseases. Front. Immunol. 12, 739430 (2021).

20. Lundtoft, C. et al. Complement C4 Copy Number Variation is Linked to SSA/Ro and SSB/La Autoantibodies in Systemic Inflammatory Autoimmune Diseases. Arthritis Rheumatol 74, 1440–1450 (2022).

21. Isenman, D. E. Chapter 17 - C4. in The Complement FactsBook (Second Edition) (eds. Barnum, S. & Schein, T.) 171–186 (Academic Press, 2018). doi:10.1016/B978-0-12-810420-0.00017-1.

22. Brandt, D. Y. C. et al. Mapping Bias Overestimates Reference Allele Frequencies at the HLA Genes in the 1000 Genomes Project Phase I Data. G3 5, 931–941 (2015).

23. Ballouz, S., Dobin, A. & Gillis, J. A. Is it time to change the reference genome? Genome Biol. 20, 159 (2019).

24. Dilthey, A. T. State-of-the-art genome inference in the human MHC. Int. J. Biochem. Cell Biol. 131, 105882 (2021).

25. Castelli, E. C. et al. HLA-G genetic diversity and evolutive aspects in worldwide populations. Sci. Rep. 11, 23070 (2021).

26. Cargou, M. et al. Evaluation of the AllType kit for HLA typing using the Ion Torrent S5 XL platform. HLA 95, 30–39 (2020).

27. Anzar, I. et al. Personalized HLA typing leads to the discovery of novel HLA alleles and tumor-specific HLA variants. HLA 99, 313–327 (2022).

28. Simakova, T. et al. NovAT tool-Reliable novel HLA alleles identification from next-generation sequencing data. HLA 99, 3–11 (2022).

29. Okada, Y. et al. Construction of a population-specific HLA imputation reference panel and its application to Graves’ disease risk in Japanese. Nat. Genet. 47, 798–802 (2015).

30. Ritari, J., et al. Increasing accuracy of HLA imputation by a population-specific reference panel in a FinnGen biobank cohort. NAR Genom Bioinform 2, lqaa030 (2020).

31. Mayor, N. P. et al. HLA Typing for the Next Generation. PLoS One 10, e0127153 (2015).

32. Houwaart, T. et al. Complete sequences of six major histocompatibility complex haplotypes, including all the major MHC class II structures. HLA 102, 28–43 (2023).

33. Luo, X., Kang, X. & Schönhuth, A. phasebook: haplotype-aware de novo assembly of diploid genomes from long reads. Genome Biol. 22, 299 (2021).

34. Ali, A. A., Aalto, M., Jonasson, J. & Osman, A. Genome-wide analyses disclose the distinctive HLA architecture and the pharmacogenetic landscape of the Somali population. Sci. Rep. 10, 5652 (2020).

35. Naslavsky, M. S. et al. Whole-genome sequencing of 1,171 elderly admixed individuals from São Paulo, Brazil. Nat. Commun. 13, 1004 (2022).

36. Silva, N. D. S. B. et al. Immunogenetics of HLA-B: SNP, allele, and haplotype diversity in populations from different continents and ancestry backgrounds. HLA 101, 634–646 (2023).

37. Abi-Rached, L. et al. Immune diversity sheds light on missing variation in worldwide genetic diversity panels. PLoS One 13, e0206512 (2018).

38. Garrison, E. et al. Variation graph toolkit improves read mapping by representing genetic variation in the reference. Nat. Biotechnol. 36, 875–879 (2018).

39. Wang, T. et al. The Human Pangenome Project: a global resource to map genomic diversity. Nature 604, 437–446 (2022).

40. Liao, W.-W. et al. A draft human pangenome reference. Nature 617, 312–324 (2023).

41. Paten, B., Novak, A. M., Eizenga, J. M. & Garrison, E. Genome graphs and the evolution of genome inference. Genome Res. 27, 665–676 (2017).

42. Rakocevic, G. et al. Fast and accurate genomic analyses using genome graphs. Nat. Genet. 51, 354–362 (2019).

43. Guarracino, A., Heumos, S., Nahnsen, S., Prins, P. & Garrison, E. ODGI: understanding pangenome graphs. Bioinformatics 38, 3319–3326 (2022).

44. Dilthey, A., Cox, C., Iqbal, Z., Nelson, M. R. & McVean, G. Improved genome inference in the MHC using a population reference graph. Nat. Genet. 47, 682–688 (2015).

45. Lee, H. & Kingsford, C. Kourami: graph-guided assembly for novel human leukocyte antigen allele discovery. Genome Biol. 19, 16 (2018).

46. Dawkins, R. et al. Genomics of the major histocompatibility complex: haplotypes, duplication, retroviruses and disease. Immunol. Rev. 167, 275–304 (1999).

47. Horton, R. et al. Variation analysis and gene annotation of eight MHC haplotypes: the MHC Haplotype Project. Immunogenetics 60, 1–18 (2008).

48. Poplin, R. et al. A universal SNP and small-indel variant caller using deep neural networks. Nat. Biotechnol. 36, 983–987 (2018).

49. Law, S. K., Dodds, A. W. & Porter, R. R. A comparison of the properties of two classes, C4A and C4B, of the human complement component C4. EMBO J. 3, 1819–1823 (1984).

50. Carroll, M. C., Fathallah, D. M., Bergamaschini, L., Alicot, E. M. & Isenman, D. E. Substitution of a single amino acid (aspartic acid for histidine) converts the functional activity of human complement C4B to C4A. Proc. Natl. Acad. Sci. U. S. A. 87, 6868–6872 (1990).

51. Dodds, A. W., Ren, X. D., Willis, A. C. & Law, S. K. The reaction mechanism of the internal thioester in the human complement component C4. Nature 379, 177–179 (1996).

52. Marin, W. M., Augusto, D. G., Wade, K. J. & Hollenbach, J. A. High-throughput complement component 4 genomic sequence analysis with C4Investigator. bioRxiv (2023) doi:10.1101/2023.07.18.549551.

53. Byrska-Bishop, M. et al. High-coverage whole-genome sequencing of the expanded 1000 Genomes Project cohort including 602 trios. Cell 185, 3426–3440.e19 (2022).

54. Mougey, R. An update on the Chido/Rodgers blood group system. Immunohematology 35, 135–138 (2019).

55. Santini, S., Boore, J. L. & Meyer, A. Evolutionary conservation of regulatory elements in vertebrate Hox gene clusters. Genome Res. 13, 1111–1122 (2003).

56. Sirén, J. et al. Pangenomics enables genotyping of known structural variants in 5202 diverse genomes. Science 374, abg8871 (2021).

57. Li, H. Aligning sequence reads, clone sequences and assembly contigs with BWA-MEM. arXiv [q-bio.GN] (2013).

58. Hickey, G. et al. Genotyping structural variants in pangenome graphs using the vg toolkit. Genome Biol. 21, 35 (2020).

59. Schneider, V. A. et al. Evaluation of GRCh38 and de novo haploid genome assemblies demonstrates the enduring quality of the reference assembly. Genome Res. 27, 849–864 (2017).

60. Norman, P. J. et al. Defining KIR and HLA Class I Genotypes at Highest Resolution via High-Throughput Sequencing. Am. J. Hum. Genet. 99, 375–391 (2016).

61. Marin, W. M. et al. High-throughput Interpretation of Killer-cell Immunoglobulin-like Receptor Short-read Sequencing Data with PING. PLoS Comput. Biol. 17, e1008904 (2021).

62. Maiers, M., Mehr, R., Raghavan, M., Kaufman, J. & Louzoun, Y. Editorial: HLA and KIR Diversity and Polymorphisms: Emerging Concepts. Front. Immunol. 12, 701398 (2021).

63. Zhang, J.-Y. et al. Using de novo assembly to identify structural variation of eight complex immune system gene regions. PLoS Comput. Biol. 17, e1009254 (2021).

64. Trowsdale, J. Genetic and functional relationships between MHC and NK receptor genes. Immunity 15, 363–374 (2001).

65. Li, H. Minimap2: pairwise alignment for nucleotide sequences. Bioinformatics 34, 3094– 3100 (2018).

66. Wick, R. R., Schultz, M. B., Zobel, J. & Holt, K. E. Bandage: interactive visualization of de novo genome assemblies. Bioinformatics 31, 3350–3352 (2015).

## References

67. Nurk, S. et al. HiCanu: accurate assembly of segmental duplications, satellites, and allelic variants from high-fidelity long reads. Genome Res. 30, 1291–1305 (2020).

68. Shafin, K. et al. Haplotype-aware variant calling with PEPPER-Margin-DeepVariant enables high accuracy in nanopore long-reads. Nat. Methods 18, 1322–1332 (2021).

69. Patterson, M. et al. WhatsHap: Weighted Haplotype Assembly for Future-Generation Sequencing Reads. J. Comput. Biol. 22, 498–509 (2015).

70. Cheng, H., Concepcion, G. T., Feng, X., Zhang, H. & Li, H. Haplotype-resolved de novo assembly using phased assembly graphs with hifiasm. Nat. Methods 18, 170–175 (2021).

71. Wang, H. & Liu, M. Complement C4, Infections, and Autoimmune Diseases. Front. Immunol. 12, 694928 (2021).

72. Excoffier, L. & Lischer, H. E. L. Arlequin suite ver 3.5: a new series of programs to perform population genetics analyses under Linux and Windows. Mol. Ecol. Resour. 10, 564–567 (2010).

73. Hickey, G. et al. Pangenome graph construction from genome alignments with Minigraph-Cactus. Nat. Biotechnol. (2023) doi:10.1038/s41587-023-01793-w.

74. Song, B. et al. AnchorWave: Sensitive alignment of genomes with high sequence diversity, extensive structural polymorphism, and whole-genome duplication. Proc. Natl. Acad. Sci. U. S. A. 119, (2022).

75. Garrison, E. & Marth, G. Haplotype-based variant detection from short-read sequencing. arXiv [q-bio.GN] (2012).

76. Mose, L. E., Perou, C. M. & Parker, J. S. Improved indel detection in DNA and RNA via realignment with ABRA2. Bioinformatics 35, 2966–2973 (2019).

77. Danecek, P. et al. Twelve years of SAMtools and BCFtools. Gigascience 10, (2021).

